# SHARP: Single-cell RNA-seq Hyper-fast and Accurate Processing via Ensemble Random Projection

**DOI:** 10.1101/461640

**Authors:** Shibiao Wan, Junil Kim, Kyoung Jae Won

## Abstract

To process large-scale single-cell RNA-sequencing (scRNA-seq) data effectively without excessive distortion during dimension reduction, we present SHARP, an ensemble random projection-based algorithm which is scalable to clustering 10 million cells. Comprehensive benchmarking tests on 17 public scRNA-seq datasets demonstrate that SHARP outperforms existing methods in terms of speed and accuracy. Particularly, for large-size datasets (>40,000 cells), SHARP’s running speed far excels other competitors while maintaining high clustering accuracy and robustness. To the best of our knowledge, SHARP is the only R-based tool that is scalable to clustering scRNA-seq data with 10 million cells.

## INTRODUCTION

By enabling transcriptomic profiling at the individual-cell level, scRNA-seq has been widely applied in various domains of biology and medicine to characterize novel cell types and detect intra-population heterogeneity (Potter 2018). The amount of scRNA-seq data in public domain has increased explosively due to technological development and the efforts to obtain large-scale transcriptomic profiling of cells (Han et al. 2018). Computational algorithms to process and analyze large-scale high-dimensional single-cell data are essential. To cluster high dimensional scRNA-seq data, dimension reduction algorithms such as principal component analysis (PCA) (Joliffe and Morgan 1992) or independent component analysis (ICA) (Hyvarinen and Oja 2000) has been successfully applied to process and to visualize high dimensional scRNA-seq data. However, it requires considerable time to obtain principal or independent components as the number of cells increases. It is also notable that dimension reduction decreases processing time at the cost of losing original cell-to-cell distances. For instance, t-distributed stochastic neighbor embedding (tSNE) (van der Maaten 2014) effectively visualizes multi-dimensional data into a reduced-dimensional space. However, tSNE distorts the distance between cells for its visualization. Besides, tSNE requires considerable time for large-scale scRNA-seq data visualization and clustering.

Random projection (RP) (Bingham and Mannila 2001) has been suggested as a powerful dimension reduction method. Based on the Johnson-Lindenstrauss lemma (Johnson and Lindenstrauss 1984), RP reduces the dimension while the distances between the points are approximately preserved (Frankl and Maehara 1988). Theoretically, RP is very fast because it does not require calculation of pairwise cell-to-cell distances or principle components.

To effectively handle very large-scale scRNA-seq data without excessive distortion of cell-to-cell distances, we developed SHARP (Supplemental Code), a hyper-fast clustering algorithm based on ensemble random projection (RP) (Fig. 1a and Methods). RP (Bingham and Mannila 2001) projects the original *D*-dimensional data into a *d*-dimensional (*d* << *D*) subspace, using a *d* × *D*-dimensional random matrix **R** whose elements conform to a distribution with zero mean and unit variance. Notably, RP preserves cell-to-cell distances even in a much lower-dimensional space and is robust to missing values, which provides a well-suited condition for clustering high-dimensional scRNA-seq data. SHARP dramatically reduced the running cost for clustering while guaranteeing far enhanced clustering performance especially for large-size scRNA-seq datasets. SHARP requires the time complexity of only 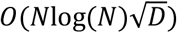 for scRNA-seq data with *N* cells and *D* genes. Compared with it, a simple hierarchical clustering algorithm requires *O*(*N*^2^*D*) (Murtagh and Legendre 2014) to calculate distance between cells. tSNE combined with the k-means algorithm requires *O*(*DN*log(*N*)) (van der Maaten 2014) and a simple PCA requires *O*(*ND* ⋅ min(*N*, *D*)) for data reduction (Bingham and Mannila 2001).

**Figure 1.**
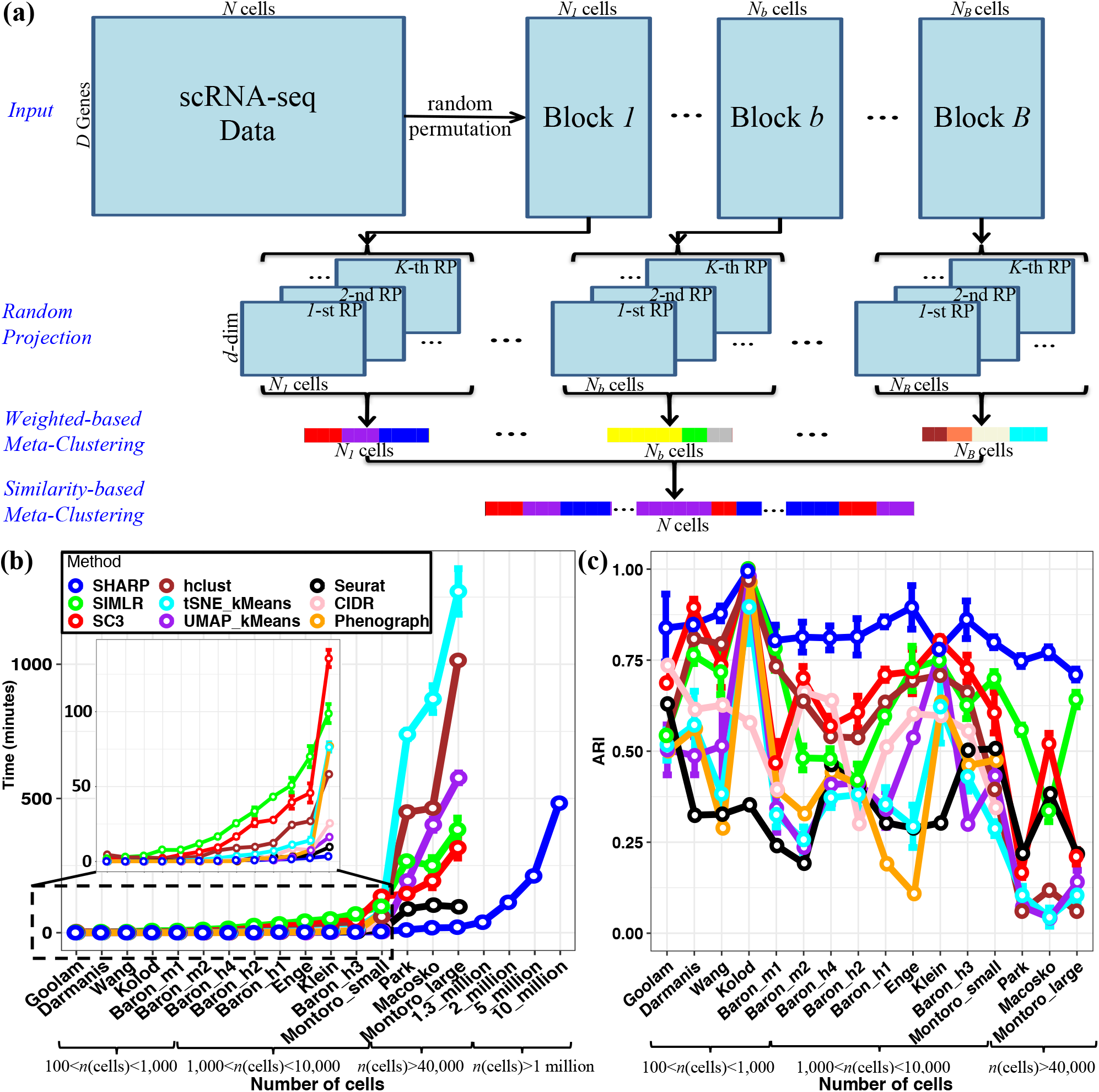
The framework of SHARP. (a) SHARP has 4 steps for clustering: divide-and-conquer, random projection (RP), weighted-based meta clustering, and similarity-based meta-clustering. (b) Running time and (c) clustering performance based on ARI (Hubert and Arabie 1985) of SHARP in 20 single-cell RNA-seq datasets with numbers of single cells ranging from 124 to 10 million (where datasets with 2 million, 5 million and 10 million cells were generated by randomly oversampling the dataset with 1.3 million single cells). For the datasets with >1 million cells, only SHARP can run and only the running time was provided due to lack of the ground-truth clustering labels. All of the results for SHARP were based on 100 runs of SHARP on each dataset. All the tests except for the larger-than-1-million-cell datasets were performed using a single core on an Intel Xeon CPU E5-2699 v4 @ 2.20GHz system with 500GB memory. To run datasets with > 1 million cells, we used 16 cores on the same system. CIDR and Phenograph were unable to produce clustering results for those datasets with number of cells larger than 40,000 (i.e., Park, Macosko and Montoro_large).

There have been previous ensemble-based approaches for RP (Fern and Brodley 2003; Bertoni and Valentini 2006). Compared with them, we developed a strategy specifically for handling scRNA-seq data by using (1) a divide-and-conquer approach for very large-scale scRNA-seq data clustering; (2) a two-layer meta-clustering approach for robust clustering; and (3) a very sparse random projection embedded into ensemble clustering for hyper-fast clustering.

We showed that SHARP preserves the cell-to-cell distance during dimension reduction and performs clustering much faster than other competitors while the clustering performance is guaranteed. We also demonstrated that SHARP is robust to the dropouts. Benchmarking various clustering algorithms, we found that SHARP is the only R-based algorithm to perform clustering 1.3 million cells (10x Genomics 2017) and can handle clustering even up to 10 million cells.

## RESULTS

### SHARP employed RP for ultra-fast scRNA-seq clustering

SHARP employed a divide-and-conquer strategy followed by RP to accommodate effective processing of large-scale scRNA-seq data (Fig. 1a and Methods). SHARP processes scRNA-seq data in 4 interconnected steps: (1) data partition, (2) RP based clustering, (3) weighted ensemble clustering and (4) similarity-based meta-clustering. During data partition, the scRNA-seq data is divided into small blocks (random size). It is noted that currently R lacks 64-bit integers support and a scRNA-seq data matrix with >1 million cells (so that the number of elements is usually significantly larger than 2^31^ − 1) cannot be directly loaded into R. The divide-and-conquer strategy enables SHARP to upload and process more than 1 million cells. The divided data blocks are further processed by RP followed by a hierarchical clustering algorithm. Because the performance of an individual RP-based clustering is volatile, ensemble of several runs of RPs is used. A weighted-ensemble clustering (i.e., wMetaC) algorithm merges individual RP-based clustering results. Finally, a similarity-based ensemble clustering (i.e., sMetaC) approach is to integrate clustering results of each block (Fig. 1a and Methods).

### SHARP is faster than other predictors and is scalable to 1.3 million cells and even up to 10 million cells

We performed comprehensive benchmarking of SHARP against existing scRNA-seq clustering algorithms including SC3 (Kiselev et al. 2017), SIMLR (Wang et al. 2017), hierarchical clustering and tSNE combined with k-means, UMAP (Becht et al. 2018) with k-means, Seurat (Butler et al. 2018), CIDR (Lin et al. 2017) and Phenograph (Levine et al. 2015) (Fig. 1b-c and Supplemental Methods) using 17 publicly available scRNA-seq datasets whose cell number ranges from 124 to 1.3 million cells (Darmanis et al. 2015; Klein et al. 2015; Kolodziejczyk et al. 2015; Macosko et al. 2015; Baron et al. 2016; Goolam et al. 2016; Wang et al. 2016; 10x Genomics 2017; Enge et al. 2017; Montoro et al. 2018; Park et al. 2018) (Supplemental Table S1).

The benchmarking tests demonstrated dramatic cost reduction by SHARP (Fig. 1b). Reflecting the theoretical running costs, the two classical algorithms (tSNE + kMeans, hierarchical clustering) manifested exponential increase in their processing time as the number of cells increased (Fig. 1b). SC3 (Kiselev et al. 2017) and SIMLR (Wang et al. 2017) showed better performance than the classical clustering approaches, but they still required a considerable amount of time for clustering. The computing cost of SHARP was substantially lower than other clustering algorithms. Remarkably, the required computing cost of SHARP rose roughly linearly even with the very large size of the datasets. For the cells with larger than 40,000, SHARP ran at least 20 times faster than SC3 (Kiselev et al. 2017) and SIMLR (Wang et al. 2017). While CIDR and Phenograph were reported to perform very fast and robustly (Levine et al. 2015; Lin et al. 2017), they were unable to produce clustering results for those datasets with more than 40,000 single cells such as Park (Park et al. 2018), Macosko (Macosko et al. 2015) and Montoro_large (Montoro et al. 2018) datasets (Fig. 1b).

Notably, SHARP clustered the scRNA-seq with 1.3 million cells in 42 minutes when using a multi-core system (Fig. 1b). Due to the data loading problem (and potential exhaustive memory use), we could not show the running time of other approaches for 1.3 million cells. When using a multi-core system (16 cores) on the Montoro_large dataset (Montoro et al. 2018) with 66,265 cells, SHARP ran remarkably faster than SC3 and SIMLR more than 40 times (Supplemental Fig S1 and Supplemental Methods). The running time of SHARP for 1.3 million cells is even 2 times (42 mins vs 96 mins) faster than that of Seurat for 66,255 cells. We expect far superior performance of SHARP against its competitors in case data loading is feasible.

To evaluate the performance of SHARP exhaustively and demonstrate the scalability of SHARP, we performed random over-sampling of the mouse brain dataset of 1.3 million cells (10x Genomics 2017) so that we were able to construct even larger sizes of scRNA-seq datasets. For this simulation, we tested up to 10 million cells. The running time of SHARP was simply linearly increased with the increasing of cell numbers from 1 million to 10 million. In our system using 16 cores, SHARP needed around 8 hours (i.e., 482.8 minutes) to cluster 10 million cells into 1175 clusters (Fig 1b).

### The outperforming clustering performance of SHARP is not highly affected by the number of cells

In parallel, we compared the clustering performance using the pre-defined cell types for each dataset (Supplemental Table S1). To evaluate clustering performance, we used adjusted Rand index (ARI) (Hubert and Arabie 1985). ARI (Supplemental Methods) is a similarity metric to measure how accurately a prediction of clustering is made in the unsupervised learning scenarios, which is similar to the accuracy measurement in supervised classification problems. Generally, the larger ARI, the better the predicted clustering is, with +1 indicating the predicted clustering is perfectly consistent with the reference, whereas 0 (or negative value) indicating that the predicted clustering is as good as (or worse than) random guess.

For almost all datasets we tested, SHARP showed better performances (Fig. 1c). It is notable that the performance of other algorithms became generally worse for large datasets (>40,000 single cells). In contrast, SHARP showed an ARI larger than 0.7 regardless of the size of the datasets, demonstrating its robustness. The ARI of Seurat was in general poor even for the small sized datasets. Seurat showed relatively faster speed (despite with worse clustering performance) even with its implementation of tSNE compared to other existing methods except SHARP. This is because Seurat only uses genes with high variations in their expressions, which could affect the clustering performance (Fig. 1c).

For robust assessment of the clustered results, we also used artificial datasets by mixing cells with known cell types obtained from Tabula Muris (Schaum et al. 2018) (Supplemental Fig S2). SHARP again outperformed other methods in terms of both clustering performance (Supplemental Fig S3) and running time (Supplemental Table S2).

### SHARP preserves cell-to-cell distance

To explain the robust clustering performance of SHARP, we investigated the degree of distortion caused by dimension reduction and compared the correlation of cell-to-cell distances after reducing dimension using SHARP, PCA and tSNE, respectively. For this, we calculated the pairwise Pearson correlation between each pairs of cells for the original scRNA-seq data and the dimension-reduced data. Reflecting the property of RP, SHARP showed almost perfect similarities in cell-to-cell distance with correlation coefficient > 0.94 even in a dimensional space which is 74 times lower (from 20862 to 279) than the original one (Fig. 2a, Supplemental Fig S4 and Supplemental Methods). Cell-to-cell distances were distorted when dimension reduction was performed to the same number of dimensions using PCA (Fig. 2a). tSNE, an algorithm to visualize high-dimensional data into 2- or 3-dimensional space, showed a lower correlation as expected (Fig. 2a).

**Figure 2.**
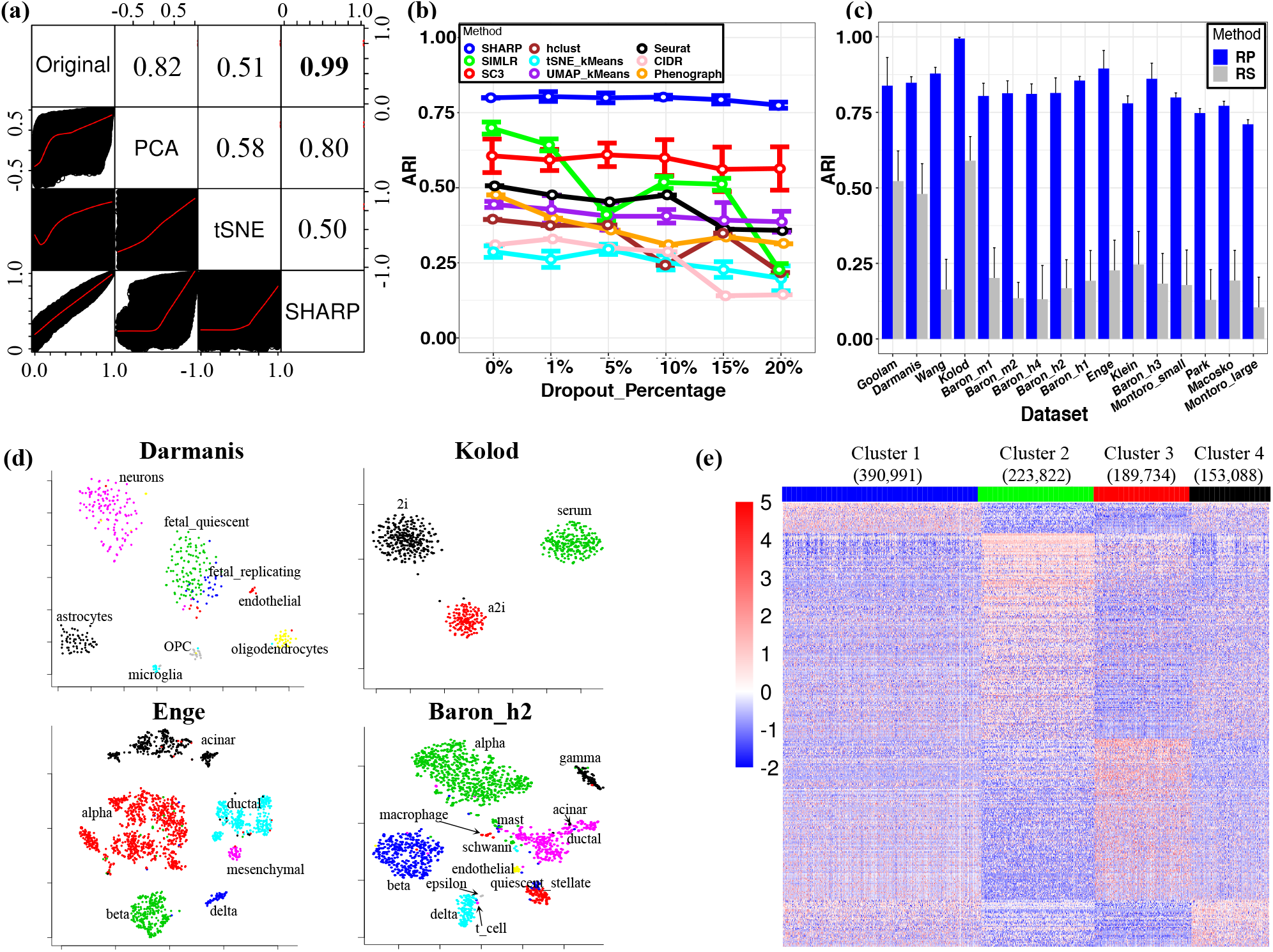
The properties of SHARP. (a) Cell-to-cell distance preservation in SHARP space comparing with that in tSNE and PCA for the Enge (Enge et al. 2017) dataset. The lower triangular part shows the scatter plots of the cell-to-cell distances, whereas the upper triangular part shows the Pearson correlation coefficient (PCC) of the corresponding two spaces. (b) SHARP is robust to the additional dropout events on the Montoro_small (Montoro et al. 2018) dataset. (c) Comparing RP (SHARP uses RP) with random gene selection (RS) in 16 single-cell RNA-seq datasets (Darmanis et al. 2015; Klein et al. 2015; Kolodziejczyk et al. 2015; Macosko et al. 2015; Baron et al. 2016; Goolam et al. 2016; Wang et al. 2016; Enge et al. 2017; Montoro et al. 2018; Park et al. 2018) with the number of single cells ranging from 124 to 66,265. (d) Visualization capabilities of SHARP in the Darmanis (Darmanis et al. 2015), Kolod (Kolodziejczyk et al. 2015), Enge (Enge et al. 2017) and Baron_h2 (Baron et al. 2016) datasets. (e) Cluster-specific marker gene expressions of the top 4 major clusters for the 1.3 million single cells (10x Genomics 2017) by SHARP. The total number of clusters predicted by SHARP is 244. The number in brackets represents the number of single cells in the corresponding cluster.

### SHARP is robust to dropouts

scRNA-seq suffers a high frequency of dropouts where many of the true expressions are not captured. To evaluate the robustness of SHARP against dropouts, we tested SHARP while increasing dropout rates in the Montoro_small (Montoro et al. 2018) dataset (Fig. 2b). It is of note that we applied additional random dropouts to the current scRNA-seq with a certain level of dropouts. For instance, originally the Montoro_small dataset has the dropout rate of 79.6% for the top 8000 genes in terms of non-zero gene expressions. Based on the original high dropout rate, additional artificial dropouts were imposed which provide more harsh conditions for clustering. Our test results showed that additional 20% of dropout was enough to evaluate the robustness of the clustering algorithms.

We found that both SHARP and SC3 were robust to the added dropouts (Methods), while we observed poorer performance for the added dropouts in general (Fig 2b). The performance of SIMLR, even though it was better than SC3 when there were no added dropouts, became worse when the added dropout rates were increased over 5%.

### Cell-to-cell distance is important for clustering results

To evaluate the contribution of RP for clustering, we performed clustering after replaying RP with random selection (RS) of genes while other procedures are unchanged. This violates the condition for RP (a matrix with zero mean and unit variance) and therefore, rough preservation of cell-to-cell distance is no longer guaranteed. Random selection of genes severely undermined the performance (Fig. 2c), suggesting that RP is a major component contributing to the performance of SHARP.

Furthermore, to demonstrate the effectiveness of wMetaC, we compared wMetaC with the method of averaging gene expressions after multiple runs of RP (we named it as “RP_avg”) while other configurations are unchanged. We observed that wMetaC performs better than simply averaging the expression profiles after random projection across all datasets (Supplemental Fig S5). The superiority of SHARP was more evident for those large-size scRNA-seq datasets when the number of cells is larger than 40,000 (e.g., Park, Macosko, and Montoro_large).

### SHARP is equipped with visualization

SHARP is equipped with tSNE (Fig. 2d and Supplemental Fig S6) and heatmaps (Supplemental Fig S7) to visualize its clustering results (Methods). For instance, the heat map for the Enge dataset (Enge et al. 2017) clearly showed the cell types in pancreas including *α* (GCG), *β* (INS), acinar (PRSS1) and *δ* (SST) cells (Supplemental Fig S7).

### Clustering 1.3 million cell data using SHARP

Of note, SHARP provides an opportunity to study the million-cell-level dataset. Previous analysis on the scRNA-seq data with 1,306,127 cells from embryonic mice brains (10x Genomics 2017) was performed using the *k*-means and a graph clustering (equivalent to kernel k-means) algorithms (10x Genomics 2017). However, *k*-means cannot identify the optimal number of clusters and it depends on the initial seeds for clustering. Using SHARP, we identified a total of 244 clusters from this dataset (17 clusters with more than 1,000 cells) (Supplemental Table S3). The top 4 clusters among them were found to have clear different expression patterns (Fig. 2e). Gene Ontology (GO) analysis (Supplemental Table S3) show that Cluster 2 is associated with dendrites and Cluster 3 is with axon. We also identified a cluster (Cluster 8) enriched for the genes associated with “non-motile cilium assembly”, which is important for brain development and function (Guemez-Gamboa et al. 2014) and immune cells with high IL4 expression (Cluster 14).

## DISCUSSION

The size of scRNA-seq data is increasing exponentially in recent years. Besides the 1.3 million brain cell dataset we used, we expect larger sizes of datasets to be generated. There is an urgent need for a computational approach to handle large size of datasets efficiently. Unfortunately, we found the majority of the computational algorithms cannot deal with very large size of the data (Fig 1b). Furthermore, the clustering performance became worse as the number of cells increases (Fig 1c).

To address these problems, we developed SHARP based on RP, which, to the best of our knowledge, has not been introduced for scRNA-seq data analysis. RP preserves the distance of the data points even in a lower dimension. Reflecting this, the cell-to-cell distance was greatly well preserved after running SHARP (Fig 2a). The performance comparison demonstrated that SHARP is a very powerful clustering software which far excels other competitors in terms of computation cost and the clustering performance (Fig 1b,c).

To further demonstrate the scalability of SHARP, we have extended our comparison of SHARP against some scalable scRNA-seq analysis methods including bigSCale (Iacono et al. 2018), Geometric Sketching (GeoSketch) (Hie et al. 2019b) and Scanorama (Hie et al. 2019a). As they were designed for different purposes, we only evaluated their scalability by running them with scRNA-seq datasets with different cell sizes and measured the running costs. Results (Supplemental Fig S8) suggested that the computing time for bigScale and Scanorama increased exponentially while SHARP and GeoSketched exerted more powerful scalability. For datasets with cell numbers larger than 200,000, bigScale and Scanorama did not work. These results demonstrate that SHARP’s scalability is comparable with other state-of-the-art algorithms.

SHARP is composed of many indispensable components. To evaluate the contribution of RP, we replaced RP with random selection, so that the Johnson-Lindenstrauss lemma (Johnson and Lindenstrauss 1984) is no longer valid. We observed the performance become severely worse by random selection, suggesting that the theoretical background by RP is a major contributor towards the performance of SHARP. Also, SHARP employed an ensemble clustering method to combine the results of several runs of RP. We found that the ensemble strategy provides the robust performance in clustering (Supplemental Fig S9 and Supplemental Methods). SHARP was robust to the ensemble size when the size is larger than 5 (Supplemental Fig S10, Supplemental Fig S11 and Supplemental Methods). Moreover, SHARP’s performance was not highly affected by the size of the block when the size is larger than 1,000 cells (Supplemental Fig S12 and Supplemental Methods). SHARP is also roughly insensitive to the degree of dimension reduction (Supplemental Fig S13 and Supplemental Methods).

It should be noted that the superior performance SHARP is not due to that a single run of RP outperforms traditional dimension reduction methods such as PCA. On the contrary, the performance of a single run of RP is volatile (Supplemental Fig S9). Instead, the superior performance of SHARP is due to the following reasons: (1) multiple runs of RP generated diversity and then they were combined with the weighted-ensemble clustering approach for robust clustering and (2) RP’s dimension reduction with the property of preserving cell-to-cell distance in the lower-dimensional space. Our meta-clustering strategies were designed to effectively handle tiny clusters generated during the data partition stage (Supplemental Fig S14 and Supplemental Note 7).

To assess the robustness of SHARP, we performed various tests. (1) We ran 100 times of SHARP and reported the mean +/− standard deviation of ARI and computational time for almost all scRNA-seq datasets in this paper; (2) We tested SHARP on a variety of scRNA-seq datasets with different numbers of cells ranging from several hundred to 10 millions; (3) We used SHARP to process scRNA-seq with various additional dropout rates. In these series of tests, SHARP demonstrated robustness across different cases.

Randomness has been widely used in multiple different domains like scalable dimensionality reduction and matrix decomposition approximation (Halko et al. 2011). The randomized method like the Gaussian random matrix, which also belongs to one kind of random projections, was used for low-rank matrix decomposition approximation (Halko et al. 2011), whereas we used random projection for dimension reduction and clustering in large-scale scRNA-seq data. Besides, the random matrix we used for SHARP was highly sparse whereas the one used in (Halko et al. 2011) was not.

Clustering involves extensive use of computational resources in calculating distances and/or dimension reduction. We demonstrated that SHARP is scalable to processing even 10 million cells. Currently, SHARP is not designed to detect rare cell populations, which could be a further application of RP. Besides clustering, the property of RP to preserve cell-to-cell distance in the reduced dimension will be useful for other applications for scRNA-seq data.

## METHODS

### The framework of SHARP

SHARP accepts gene expression data arranged in a matrix 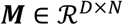, where each of the *D* rows corresponds to a gene (or transcript) and each of the *N* columns corresponds to a single cell. The type of input data can be either fragments/reads per kilo base per million mapped reads (FPKM/RPKM), counts per million mapped reads (CPM), transcripts per million (TPM) or unique molecule identifiers (UMI). For consistency, FPKM/RPKM values are converted into TPM values and UMI values are converted into CPM values.

### Data partition

For a large-scale dataset SHARP performs data partition using a divide-and-conquer strategy. SHARP divides scRNA-seq data 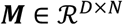 into *B* blocks, where each block may contain different numbers of cells (i.e., *N*_1_, …, *N*_B_, where 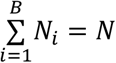). To avoid bias during data partition, we randomly permuted the original single-cell data before partitioning. In practice, SHARP roughly equally divides ***M*** and allows users to assign the base number of single cells in each block (e.g., *n*). In this case, *B* = ⌈*N*/*n*⌉, where ⌈*x*⌉ is the minimum integer no less than *x*. The numbers of single cells 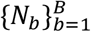 in each block are as follows:

1. If *B* = 1, *N*_*b*_ = *N*, where *b* = *B* = 1;
2. If *B* = 2, 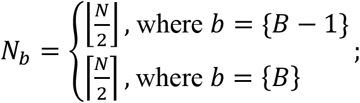
3. If *B* ≥ 3, 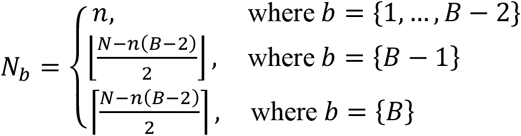

This enables SHARP to maximize the usage of local computational resources and avoid memory overflow while minimizing the negative impact from imbalanced numbers of data for each block.

### Random Projection (RP)

RP is a group of simple yet powerful dimension-reduction technique. It is based on the Johnson-Lindenstrauss lemma (Johnson and Lindenstrauss 1984) (Supplemental Methods). Specifically, the original *D*-dimensional data is projected onto a *d*-dimensional subspace, using a random matrix whose column are unit length, i.e.,

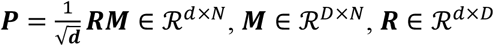

As long as the elements of **R** conforms to any distributions with zero mean and unit variance, **R** gives a mapping that satisfies the Johnson-Lindenstrauss lemma.

### Choice of random matrix R

To reduce the computational complexity, we adopted a very sparse RP proposed in (Li et al. 2006) where the elements of **R** (i.e., *r*_*i,j*_) is defined as:

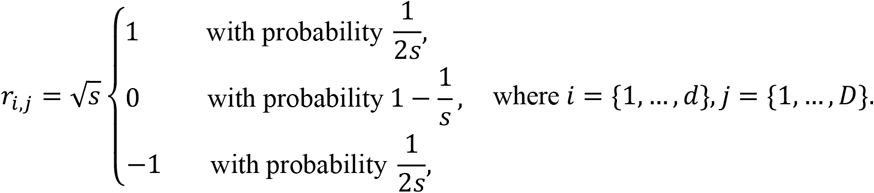

As suggested in (Li et al. 2006), we selected 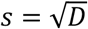.

### Choice of the subspace dimension *d*

To balance between maintaining robust performance and yielding a solution as parsimonious as possible, we selected *d* = log_2_(*N*)/*ϵ*^2^, where *ϵ* ∈ (0,1] as suggested by Johnson-Lindenstrauss lemma.

### Ensemble RP

After RP, pairwise correlation coefficients between each pair of single cells were calculated using the dimension-reduced feature matrix. An agglomerative hierarchical clustering (hclust) with the “ward.D” (Ward Jr 1963) method was used to cluster the correlation-based distance matrix. We first applied RP *K* times to obtain *K* RP-based dimension-reduced feature matrices and then further *K* distance matrices. Each of the K matrices was clustered by a “ward.D”-based hclust. As a result, *K* different clustering results were obtained, each from a RP-based distance matrix, which would be combined by a weighted-based meta-clustering (wMetaC) algorithm (Ren et al. 2017) detailed in the next step.

### Weighted-based meta-clustering (wMetaC)

Compared to the traditional cluster-based similarity partitioning algorithm (CSPA) (Strehl and Ghosh 2002) which treat each instance and each cluster equally important, wMetaC assigns different weights to different instances (or instance pairs) and different clusters to improve the clustering performance. wMetaC includes four steps: (1) calculating cell weights; (2) calculating weighted cluster-to-cluster pairwise similarity; (3) clustering on weighted cluster-based similarity matrix and (4) determining final results by a voting scheme. Note that wMetaC was applied to each block of single cells. The flowchart of the wMeaC ensemble clustering method is shown in Supplemental Fig S15. Specifically, for calculating cell weights, similar to the first several steps in CSPA, we first converted the individual RP-based clustering results into a co-location similarity matrix **S**, whose element *s*_*i,j*_ represents the similarity between the *i*-th and *j*-th single cells. Then, based on the idea that the weight for each pair of single cells is determined by the degree of consistency of the co-location clustering results of these two single cells, we converted the similarity matrix **S** to the weight matrix **W** according to the following equation:

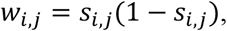

where *w*_*i,j*_ is the element in the *i*-th row and the *j*-th column of W. It is easy to see that when *s*_*i,j*_ = 1 (i.e., the *i*-th cell and the *j*-th cell are with 100% probability in the same cluster) or *s*_*i,j*_ = 0 (i.e., the *i*-th cell and the *j*-th cell are with 0% probability in the same cluster), *w*_*i,j*_ reaches the minimum at 0; when *s*_*i,j*_ = 0.5 (i.e., the co-location probability of the *i*-th cell and the *j*-th cell in the same cluster is 0.5 whereas the probability of them in different clusters is also 0.5, which means this is the most difficult-to-cluster case), *w*_*i,j*_ reaches the maximum at 0.25. In other words, zero weight is assigned to those most “easy-to-cluster” pairs of single cells and the highest cell-to-cell weight is assigned for the most “difficult-to-cluster” pairs. Then, a weight associated with each cell was calculated as the accumulation of all the cell-to-cell weights related with the corresponding cell. To calculate the weighted cluster-to-cluster similarity, we first noted that the size of the similarity matrix is |*C*| × |*C*|, where *C* is the union set of all the clusters obtained in each individual RP-based clustering results in the previous step, and | ⋅ | is the cardinality of a set. Then, for any two clusters, their similarity is determined by the sum of weights of their overlapped elements (i.e., cells) divided by that of their combined ones. Specifically, given two clusters *C*_*u*_ and *C*_*ν*_, the cluster-to-cluster similarity in wMetaC is defined as:

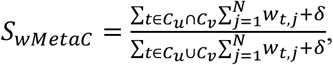

where *w*_*t,j*_ is the co-location weight for the *t*-th cell and *j*-th cell derived above, *N* is the number of cells, and *δ* is a very small positive number (by default, we used *δ* = 0.01) to avoid the denominator to be 0. We can treat 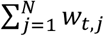 as the overall co-location weights for the *t*-th cell because it sums up all the possible colocation weights between the *t*-th cell and all cells. Thus, in the equation, the numerator represents the sum of the co-location weights of the cells which are found in both Cluster *C*_*u*_ and Cluster *C*_*ν*_, whereas the denominator represents the sum of the co-location weights of the cells which are found in either Cluster *C*_*u*_ or Cluster *C*_*ν*_.

Then, in the third step (i.e., meta-clustering), we used a hierarchical clustering with “ward.D” to cluster the obtained similarity matrix. After clustering, we understood which cluster in the 1-st RP-based clustering corresponds to which cluster(s) in the 2-nd, 3-rd,…, *K*-th RP-based clustering. Then, in the final step, we reorganized the *K* RP-based clustering results according to the result in the third step, and then we used a voting scheme (see below) to determine the final clustering results. These procedures were repeated for each of the *B* blocks.

### Voting scheme

Given *K* runs of RPs, we obtained *K* different individual clustering results by using hierarchical clustering on each of the *K* RP-based dimension-reduced feature matrices. Suppose that the *k*-th (*k* ∈ {1, …, *K*}) individual clustering results in |*C*^*k*^| clusters, i.e., 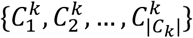. Then, by using the first three steps of wMetaC, we obtained a weighted-based cluster-wise similarity matrix and then we used hierarchical clustering again for this meta-clustering problem. After meta-clustering, assume that the number of clusters is |*G*|, i.e., {*G*_1_, …, *G*_*g*_, …, *G*_|*G*|_}, where 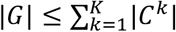. Thus, *G*_*g*_ corresponds to one or more clusters from the individual clustering set. Finally, a single cell was assigned to the meta-cluster to which it belongs with the highest ratio. Ties were broken randomly. For example, given *K* = 5, suppose for a single cell, its 5 RP-based individual clustering results are 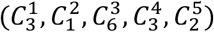, i.e., it belongs to the 3-rd cluster in the 1-st RP clustering results, the 1-st cluster in the 2-nd RP, the 6-th cluster in the 3-rd RP, the 3-rd cluster in the 4-th RP and the 2-nd cluster in the 5-th RP (Note that the individual clusters are not necessarily consistent with each other and that’s why meta-clustering method like wMetaC is required). After meta-clustering, suppose that 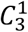, 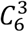 and 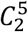 belong to the same meta-cluster *G*_3_, while 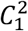 and 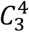 belong to *G*_1_ and *G*_2_, respectively. Because this single cell is predicted to belong to Cluster *G*_3_ with the highest ratio (i.e., 3/5), the final predicted cluster of this cell is Cluster *G*_3_.

### Similarity-based meta-clustering (sMetaC)

To integrate the clustering results of the *B* blocks obtained by wMetaC, we proposed a similarity-based meta-clustering (sMetaC) approach which is similar to wMetaC. The major differences between wMetaC and sMetaC: (1) the cluster-to-cluster pairwise similarity of the former is calculated based on co-location weights of single cells in each cluster, whereas that of the latter is calculated based on the mean of the cell-to-cell correlation coefficients; (2) the individual clustering results of the former actually correspond to the same block of single cells but in different lower-dimensional space, whereas those of the latter correspond to different blocks of single cells; (3) the former requires a voting scheme to integrate K individual clustering results, whereas the latter does not and it just needs to reorganize the clusters to make clusters consistent across blocks.

The cluster-to-cluster pairwise similarity of wMetaC is calculated based on co-location weights of single cells in each cluster, whereas that of the sMetaC is calculated based on the Pearson correlation coefficient of the mean cluster-wise feature vectors after ensemble random projection. Specifically, in sMetaC, given two clusters *G*^*u*^ and *G*^*ν*^ which were obtained by wMetaC (note that these two clusters may or may not belong to the same block of single cells), we calculated the similarity between these two clusters in sMetaC as follows

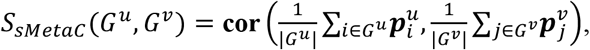

where **cor**(⋅,⋅) is the Pearson correlation coefficient of two vectors, 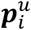 and 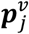 are the dimension-reduced feature vectors after ensemble random projection for the *i*-th cell in Cluster *G*^*u*^ and the *j*-th cell in Cluster *G*^*ν*^, and |*G*^*u*^| and |*G*^*ν*^| are the numbers of cells in Cluster *G*^*u*^ and Cluster *G*^*ν*^, respectively.

### Determining the optimal number of clusters

SHARP determines the optimal number of clusters by using three criteria which are based on internal evaluations of the clustering results (Supplemental Methods).

### Time complexity analysis

SHARP includes 4 steps for clustering: (1) data partition; (2) RP; (3) wMetaC; (4) sMetaC. For a scRNA-seq data matrix 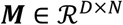, SHARP first divides the data into *B* blocks, the *b*-th block with *N*_*b*_ single cells. According to the “data partition” section, *N*_*b*_ ≤ *n*, where *n* is a fixed user-defined parameter enabling that one application of RP-based clustering runs sufficiently fast. Our analysis (Supplemental Fig S12) shows that *n* = 1500 or 2000 is a good balance between performance and speed in our case. Then, for each block, one run of RP requires time complexity of 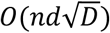 (Johnson and Lindenstrauss 1984), where *d* = ⌈log(*N*)/*ϵ*^2^⌉ ≪ *D*. Note here, *d* is calculated based on *N* rather than *n* for dimension-reduction consistency across blocks. Practically, in the 13 reported scRNA-seq datasets, the dimension can be reduced by 42~238 times (i.e., *D/d*), depending on the number of single cells and the number of genes. SHARP requires several (i.e., *K*) runs of RPs (with complexity of 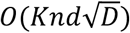. Subsequently, SHARP uses a hierarchical clustering (hclust) with “ward.D” for each of the *K* RPs, thus with overall time complexity of 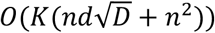 (Note that the time complexity of hclust in R package is *O*(*n^2^*) (Murtagh and Legendre 2014)). Later, wMetaC, essentially a hclust for the individual predicted clusters (also without loss of generality, suppose the number of clusters for each RP-based clustering is equally *C*_2_, where *C*_2_ ≪ *n*), was applied to each block (in total, time complexity of 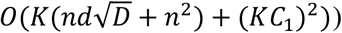. Finally, SHARP integrated the results of all blocks by proposing a method called sMetaC, whose time complexity is similar to wMetaC except the number of instances is different (similarly, we can suppose the number of clusters in each block is equally *C*,, where *C*, ≪ *n*). In this case, the total time complexity is 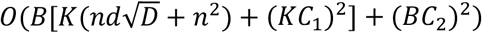. Practically, *K, B*, *C*_1_ and *C*_2_ are very small, thus the time complexity of SHARP can be written as 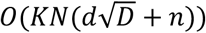. Because *n* is fixed across different datasets, *d* = ⌈log(*N*)/*ϵ*,⌉ and D is usually larger than 10,000, thus 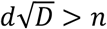 and therefore the time complexity of SHARP is essentially 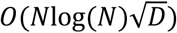.

On the other hand, among the compared state-of-the-art methods, tSNE plus k-means is arguably the fastest. Theoretically, tSNE requires *O*(*DN*log(*N*)) for dimension reduction to 2D or 3D space (van der Maaten 2014). For tSNE plus k-means for clustering, the time complexity is *O*(*DN*log(*N*) + 2*Nki*), where *k* is the number of clusters and *i* is the number of iterations. Thus, the total time complexity for tSNE + kMeans is *O*(*DN*log(*N*)).

All the tests except for the 1.3-million-cell dataset were performed using a single core on an Intel Xeon CPU E5-2699 v4 @ 2.20GHz system with 500GB memory. To run 1.3 million cells, we used 16 cores on the same system.

### Visualization

For visualization, SHARP uses a weighted combination of the dimension-reduced feature matrix and the cell-to-cluster matrix derived from the clustering results. The former matrix is obtained by three steps: 1) applying *K* runs of RPs for each block of the large-scale scRNA-seq data; 2) combining these block-wise matrices to obtain *K* RP-based dimension-reduced matrices and 3) averaging these *K* matrices into one ensemble matrix. For the latter matrix, we constructed a *N* × *pC* matrix, where *N* is the number of single cells and *pC* is the predicted number of clusters. If the *i*-the single cell is predicted to be in the *j*-th cluster, then the element of the *i*-th row and *j*-th column is 1, otherwise 0. Subsequently, these two matrices were combined with different weights to formulate the visualization matrix, which is the input matrix of tSNE for visualization.

First, both the dimension-reduced feature matrix 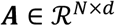 (where *N* is the number of cells and *d* is the number of dimensions after dimension reduction) and the cell-to-cluster matrix 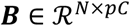 (where *pC* is the predicted number of clusters) are centered and scaled along each dimension across cells, and we notate the results as 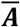 and 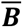. Then, these two matrices are combined as follows: 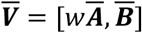, where 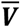; is the input matrix to tSNE for visualization and *w* >0 is the weight ratio of the dimension-reduction feature matrix over the cell-to-cluster matrix. If 0<*w*<1, more weight will be given to cell-to-cluster matrix, suggesting that the clustering results are believed to be better for visualization; if *w*>1, more weight will be given to the dimension-reduced feature matrix, which indicates that the data in dimension-reduced space can be more suitable for visualization. While it is possible to employ some algorithms to optimize *w*, we adopted an empirical value (i.e., *w* = 2) by default, which is robust for better visualization across different scRNA-seq datasets. For flexibility, we also provided an extra option to allow users to define by their own *w*.

Based on the clustering results, SHARP can further detect genes for each cluster. We adopted a method similar to SC3 except three points: (1) Besides p-value and area under receiver operating curve (AUROC), SHARP uses two more criteria to select marker genes, namely cluster-mean fold change (FC) and expression sparsity (i.e., the percentage of expressions across all cells); (2) SHARP uses an adaptive threshold instead of a hard-threshold (i.e., p-value < 0.01 and AUROC > 0.85); (3) SHARP uses a parallelization way to calculate all of the criteria mentioned above.

### Simulated data

In this study, we have generated three simulated scRNA-seq data from the Tabula Muris (Schaum et al. 2018). Specifically, we only selected the data from the microfluidic droplet-based method and three simulated data were generated, i.e., mdata3, mdata6 and mdata8. mdata3 was generated by mixing scRNA-seq data from three organs, including heart and aorta, thymus and liver; mdata6 was generated from six organs, including heart and aorta, thymus, liver, bladder, kidney and tongue; and mdata8 was generated from eight organs, including, heart and aorta, thymus, liver, bladder, kidney, tongue, spleen and trachea. Note that only those filtered data by the original paper were used. The total numbers of single cells for mdata3, mdata6 and mdata8 are 3,898, 16,717 and 37,538, respectively.

### Adding dropouts

To further demonstrate the robustness of SHARP against scRNA-seq dropout events, we artificially added dropouts to a benchmarking dataset (e.g., Montoro_small). Specifically, we randomly selected a percentage (e.g., 1%, 5%, 10%, 15% and 20%) of non-zero expressions from the Montoro_small dataset and then set them to be 0. In other words, the dropout percentage here refers to the added dropout percentage.

#### Public datasets used in this study

Single cell RNA-seq data were obtained from the GEO accession numbers provided by their respective original publications. For the 1.3 million single-cell dataset, we downloaded from the 10x Genomics website https://support.10xgenomics.com/single-cell-gene-expression/datasets/1.3.0/1M_neurons. The scRNA-seq data from the Tabula Muris was downloaded from the website https://figshare.com/projects/Tabula_Muris_Transcriptomic_characterization_of_20_organs_and_tissues_from_Mus_musculus_at_single_cell_resolution/27733.

## Software Availability

The source code for SHARP has been submitted to Github (https://github.com) under the project name SHARP (https://github.com/shibiaowan/SHARP).

## ACKNOWLEDGEMENTS

This work is supported by NIH grant (R01 DK106027) and the Novo Nordisk Foundation grant number NNF17CC0027852.

## AUTHORS’ CONTRIBUTIONS

SW and KJW conceived and designed the study. SW developed the SHARP algorithm, performed the experiments and analyzed the data. SW implemented the SHARP package. SW, JK and KJW participated in writing the paper. The manuscript was approved by all authors.

