## Supplemental Fig S1 for "SHARP: Single-cell RNA-seq Hyper-fast and Accurate Processing via Ensemble Random Projection"

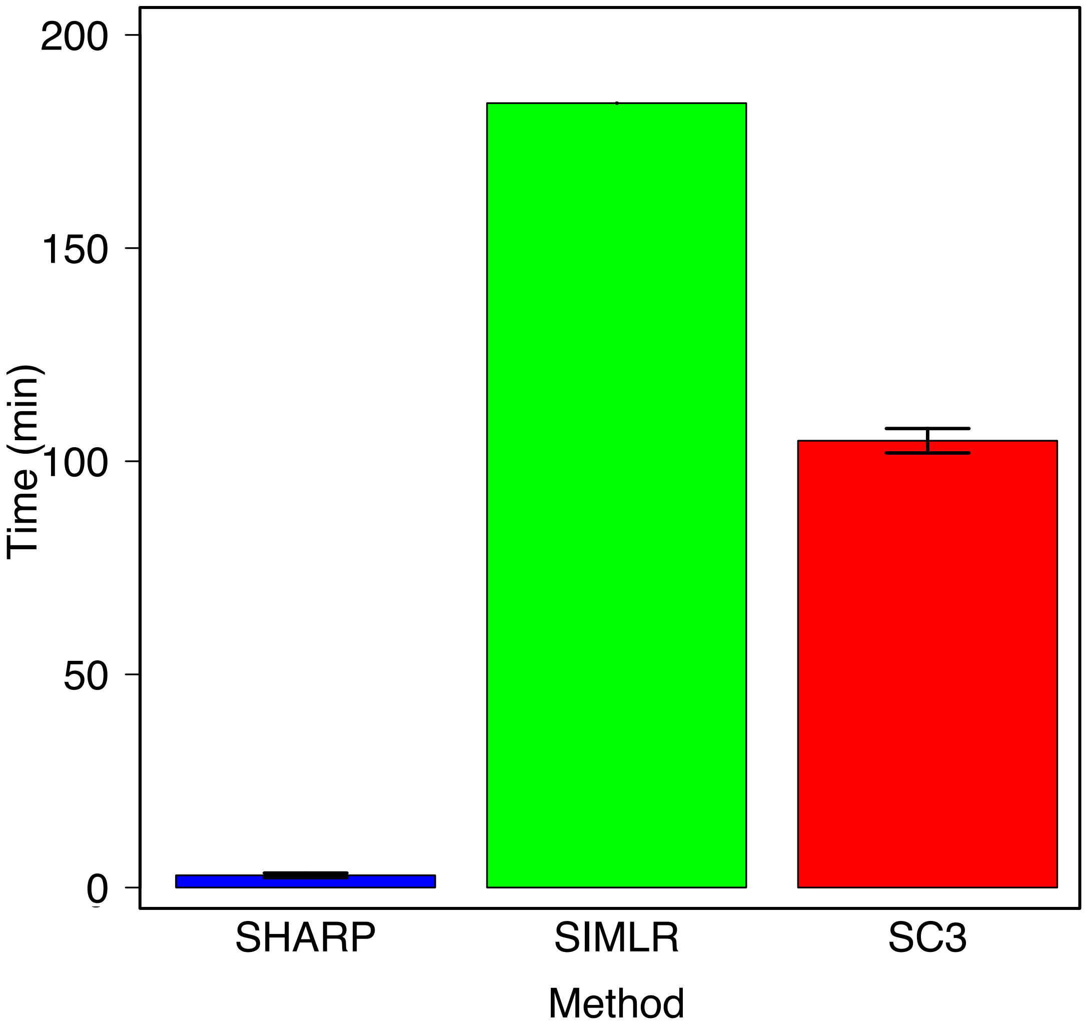


**Supplemental Fig S1**: Comparison of running cost for SHARP, SC3 and SIMLR on the Montoro et al. dataset [21] (containing 66,265 single cells) using multi-core configurations (16 cores). The results for SHARP and SC3 are based on 100 runs of the corresponding algorithms.
