## Supplemental Fig S2 for "SHARP: Single-cell RNA-seq Hyper-fast and Accurate Processing via Ensemble Random Projection"

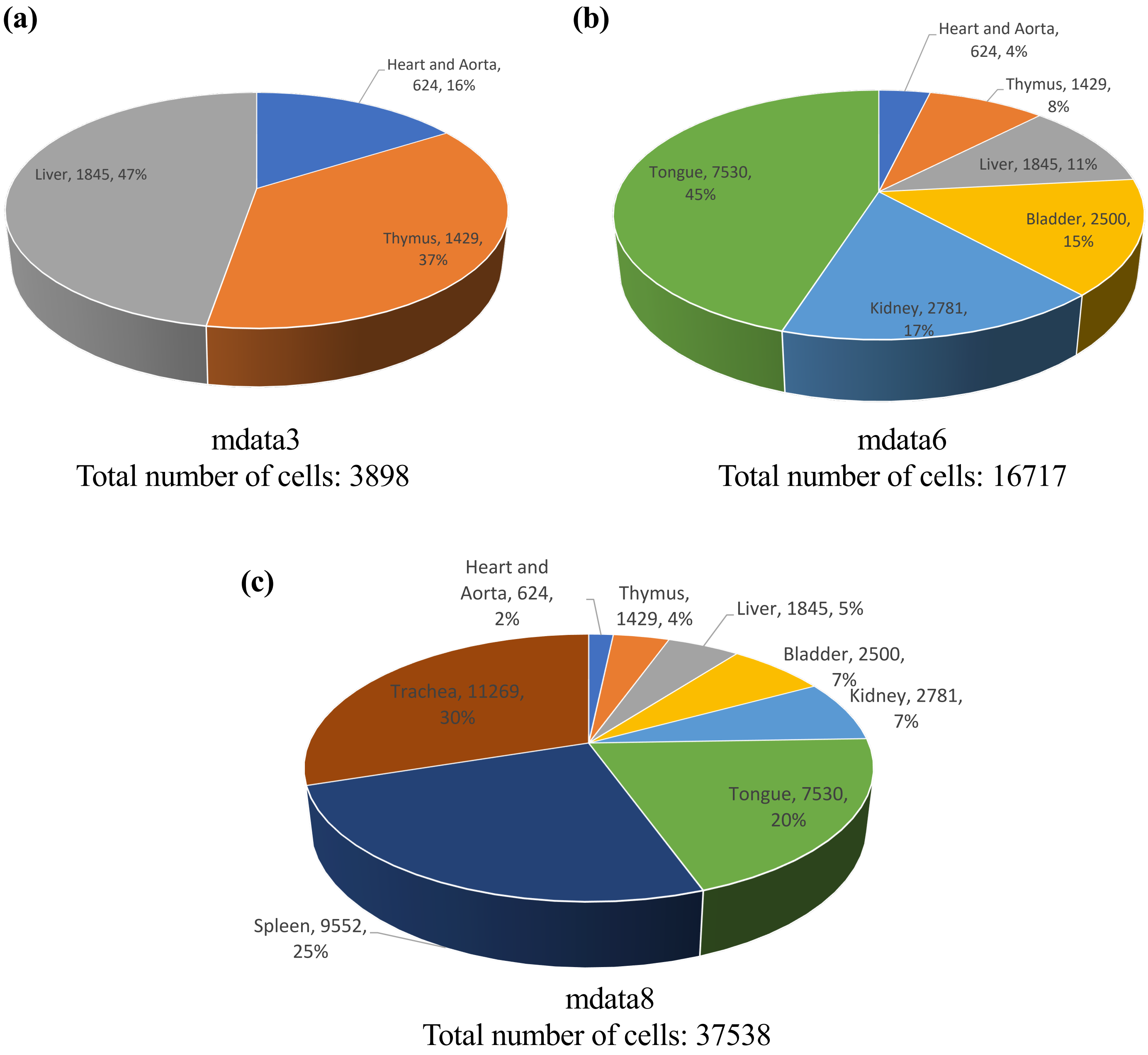


**Supplemental Fig S2**: Breakdown of the three simulated scRNA-seq data from the Tabula Muris [26]. (a) mdata3 consists of data from three organs, including heart and aorta, thymus and liver. (b) mdata6 consists of data from six organs, including heart and aorta, thymus, liver, bladder, kidney and tongue. (c) mdata8 consists of data from eight organs, including heart and aorta, thymus, liver, bladder, kidney, tongue, spleen and trachea. The total numbers of cells for mdata3, mdata6 and mdata8 are 3,898, 16,717 and 37,538, respectively.
