## Supplemental Fig S3 for "SHARP: Single-cell RNA-seq Hyper-fast and Accurate Processing via Ensemble Random Projection"

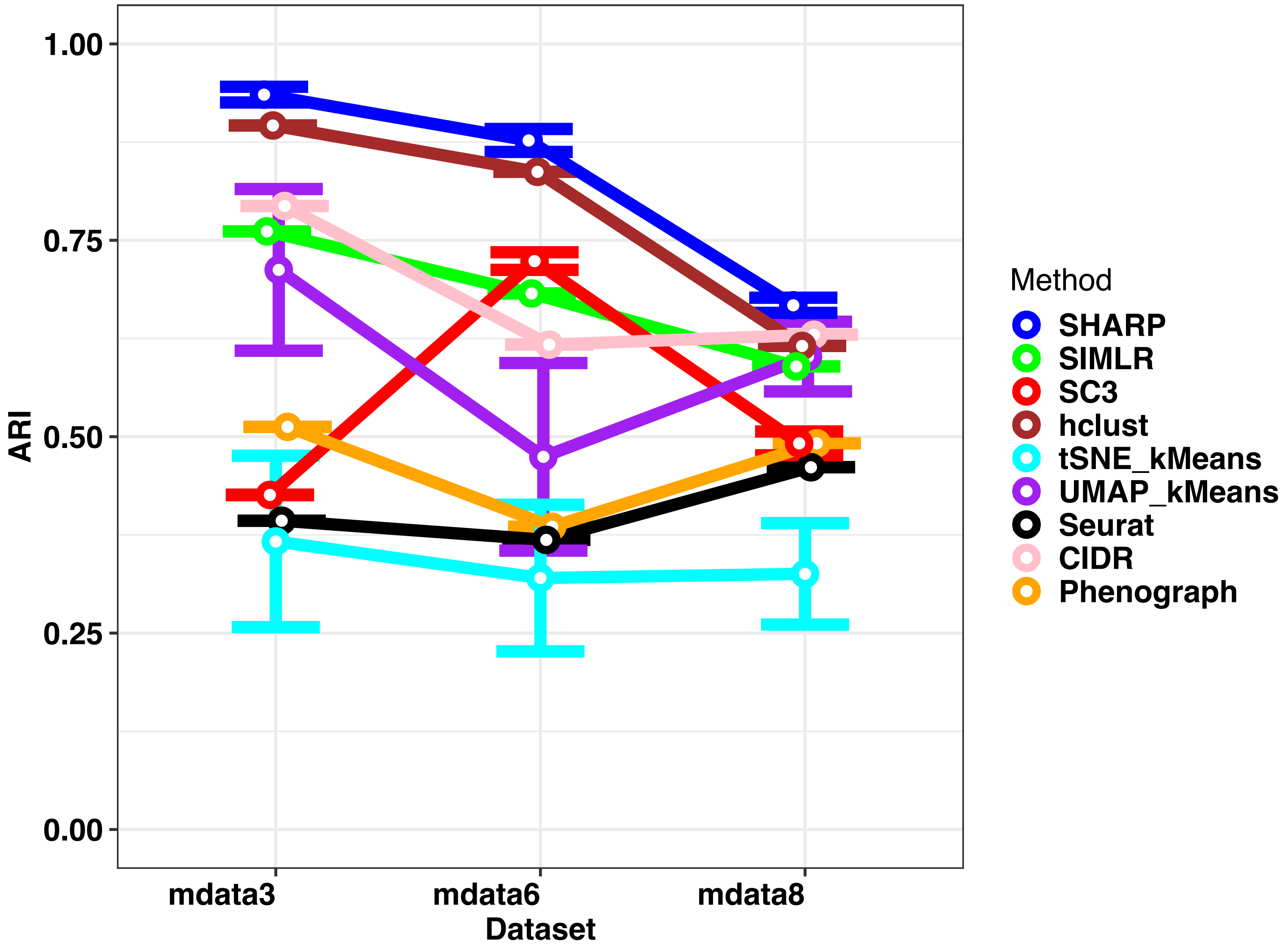


**Supplemental Fig S3**: Comparing SHARP with existing methods on simulated scRNA-seq datasets from the Tabula Muris [26] by using 3, 6 and 8 organs, respectively. mdata3: mixing data from three organs, including heart and aorta, thymus and liver; mdata6: mixing data from six organs, including heart and aorta, thymus, liver, bladder, kidney and tongue; and mdata8: mixing data from eight organs, including, heart and aorta, thymus, liver, bladder, kidney, tongue, spleen and trachea. The total numbers of single cells for mdata3, mdata6 and mdata8 are 3,898, 16,717 and 37,538, respectively.
