## Supplemental Fig S4 for "SHARP: Single-cell RNA-seq Hyper-fast and Accurate Processing via Ensemble Random Projection"

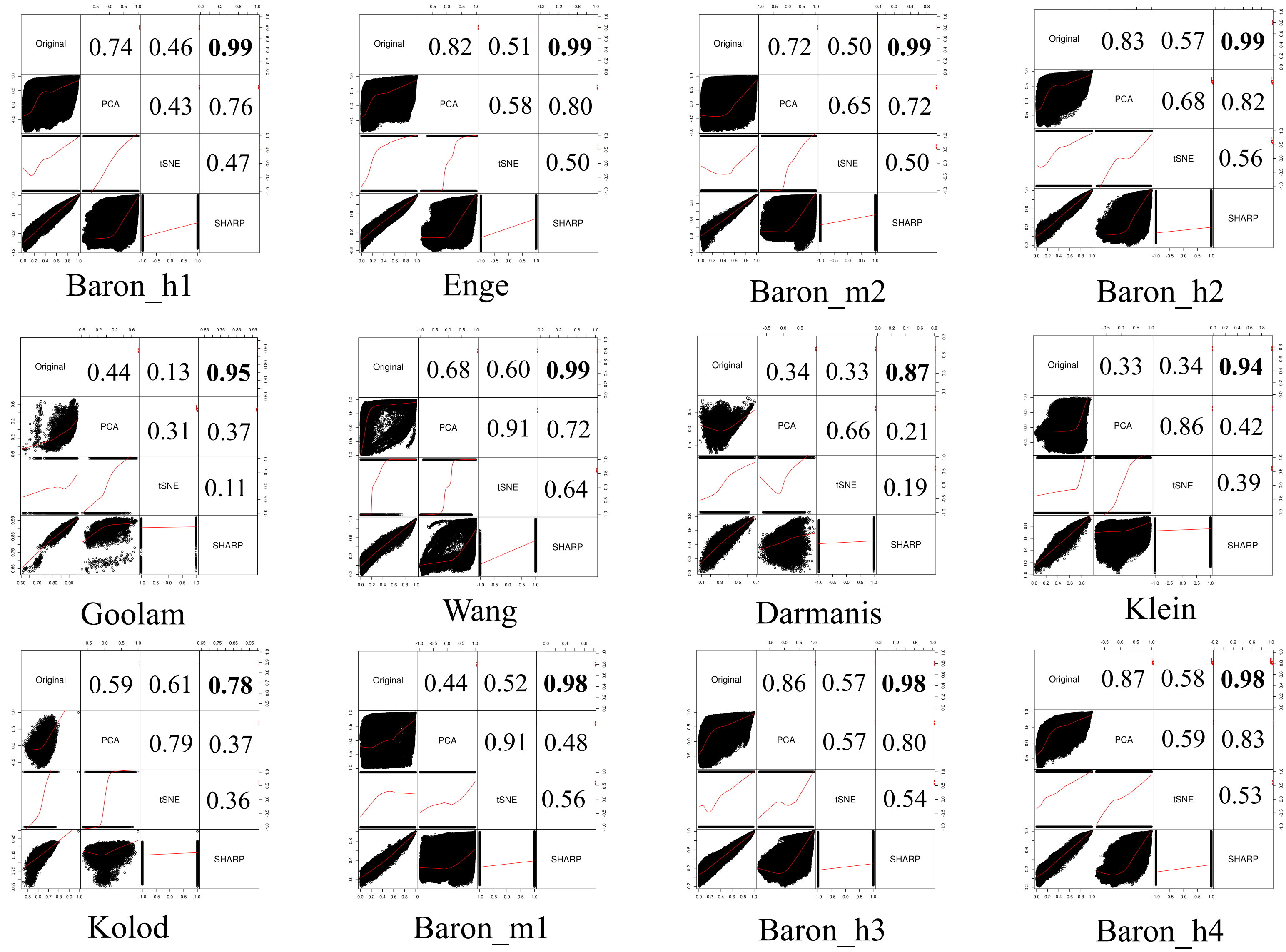


**Supplemental Fig S4**: Cell-to-cell distance preservation in SHARP. The lower triangular part shows the scatter plots of the cell-to-cell distances and the upper triangular part shows the Pearson correlation coefficient (PCC) of the corresponding two spaces. The dimension for SHARP and PCA is reduced to $\log_2(N)/0.2^2 = 25\log_2(N)$ from the original dimension $D$, where $N$ and $D$ are the numbers of cells and genes, respectively. The dimension of tSNE is 3.
