## Supplemental Fig S5 for "SHARP: Single-cell RNA-seq Hyper-fast and Accurate Processing via Ensemble Random Projection"

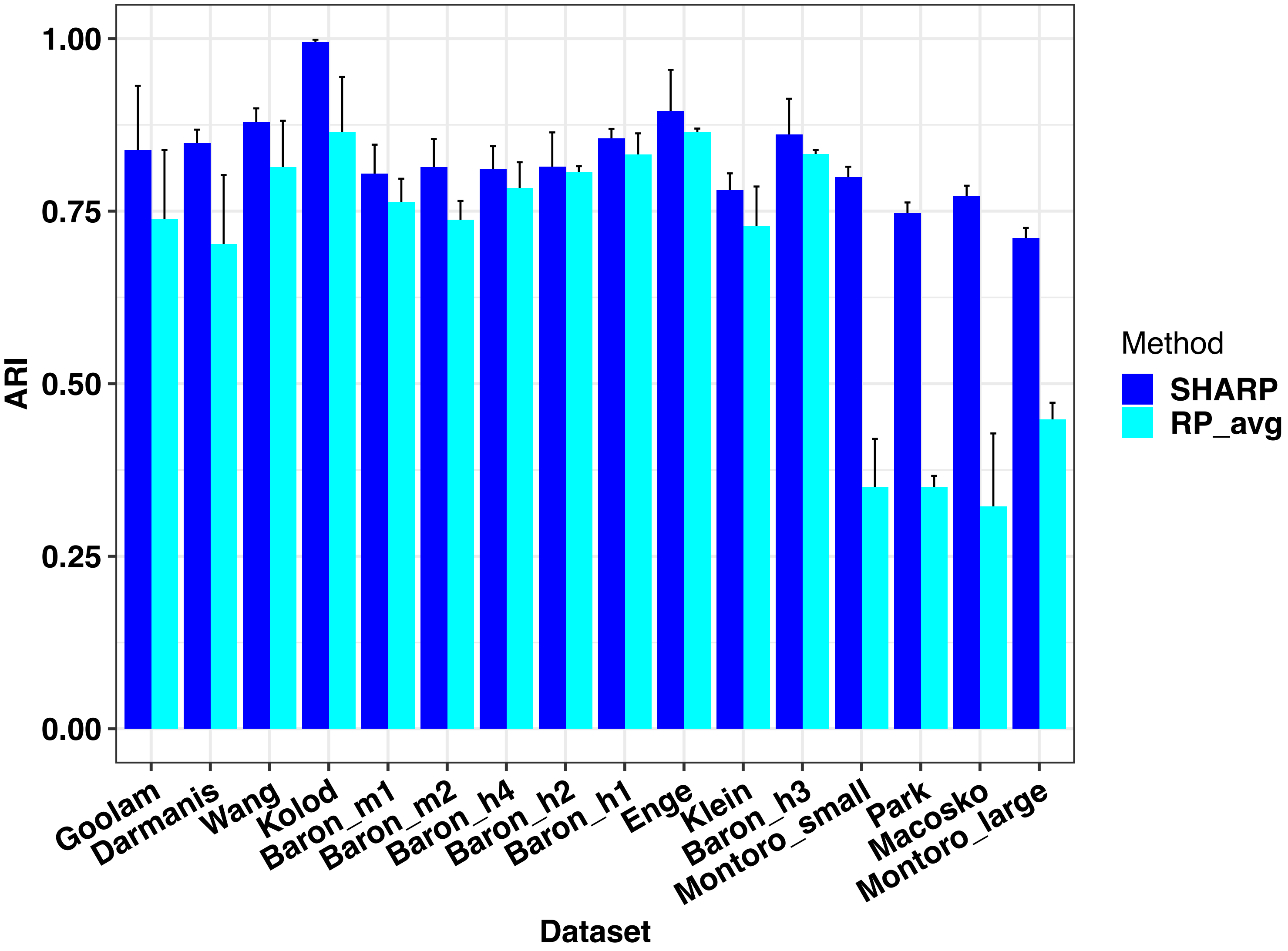


**Supplemental Fig S5**: Comparing wMetaC with the method of averaging gene expressions after multiple runs of RP (i.e., "RP_avg"). We used the same configurations (e.g., random permutation, random projection, sMetaC, etc) for RP_avg as SHARP except that wMetaC was replaced by simply averaging gene expressions.
