## Supplemental Fig S6 for "SHARP: Single-cell RNA-seq Hyper-fast and Accurate Processing via Ensemble Random Projection"

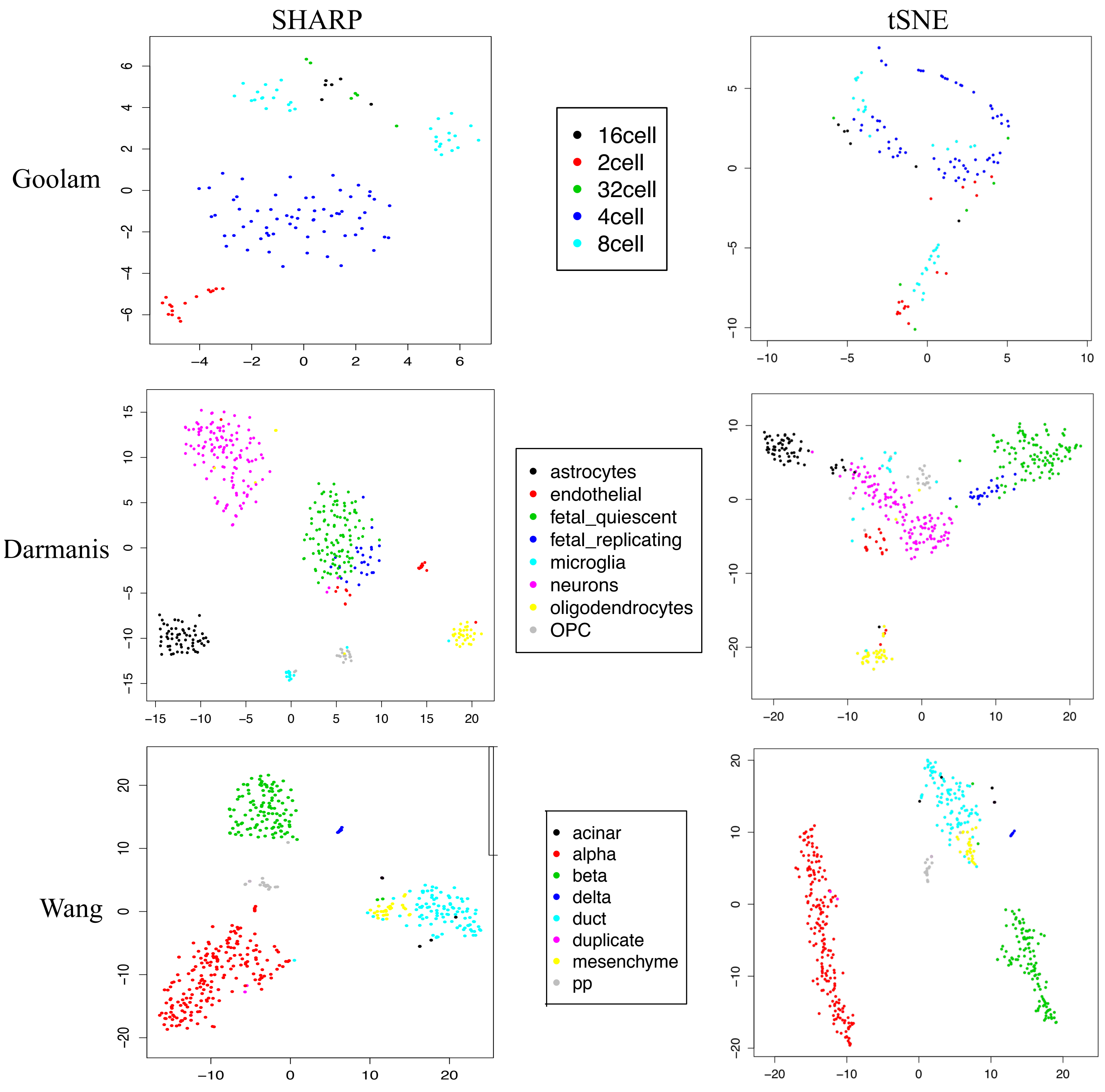


**Supplemental Fig S6a**: Visualization capabilities of SHARP compared with that of tSNE in the Goolam [14], Darmanis [15] and Wang [16] datasets. The numbers of single cells in these three datasets are 124, 420 and 479, respectively.

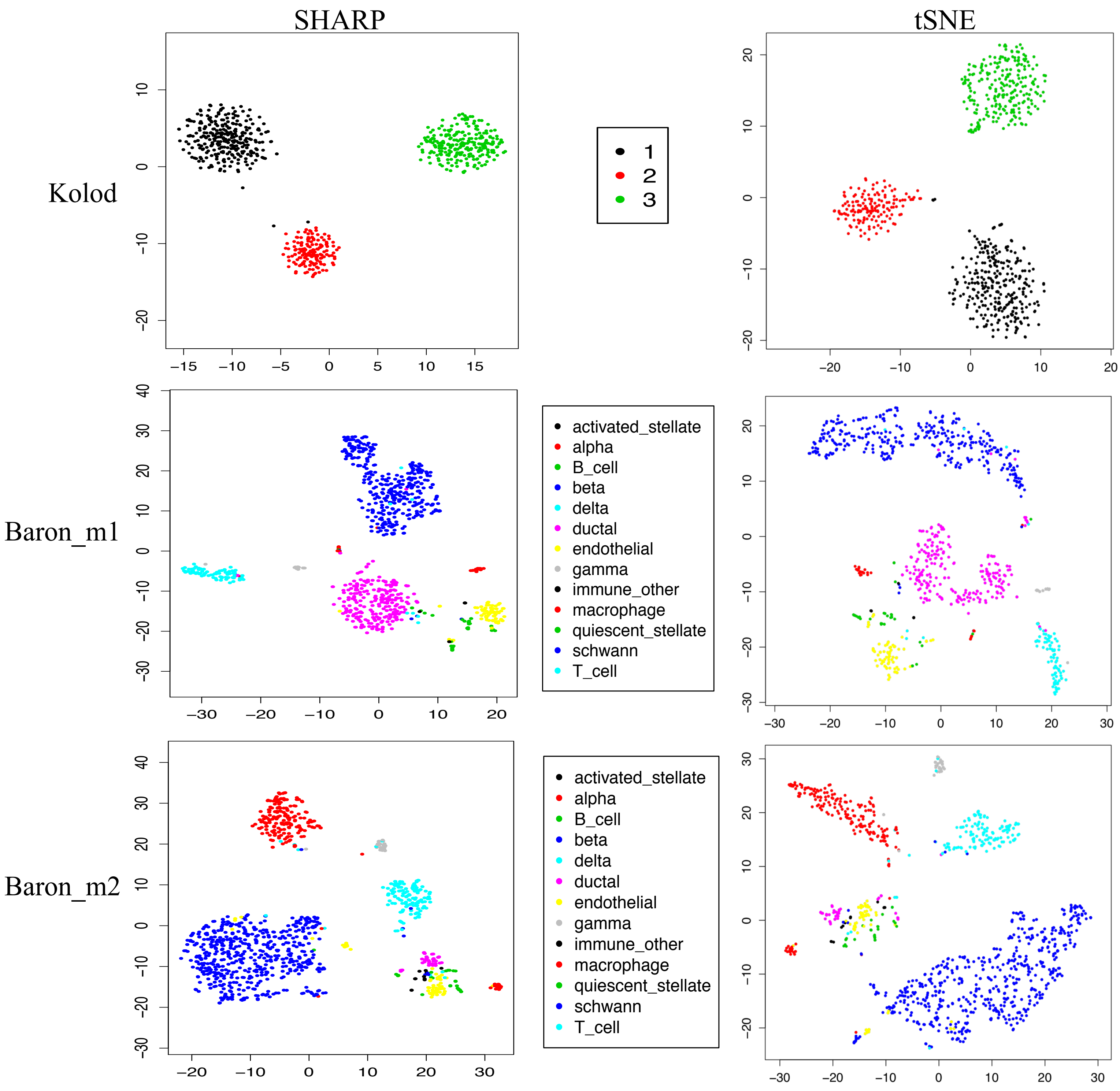


**Supplemental Fig S6b**: Visualization capabilities of SHARP compared with that of tSNE in the Kolod [17], Baron_m1 [18] and Baron_m2 [18] datasets. The numbers of single cells in these three datasets are 704, 822 and 1064, respectively.

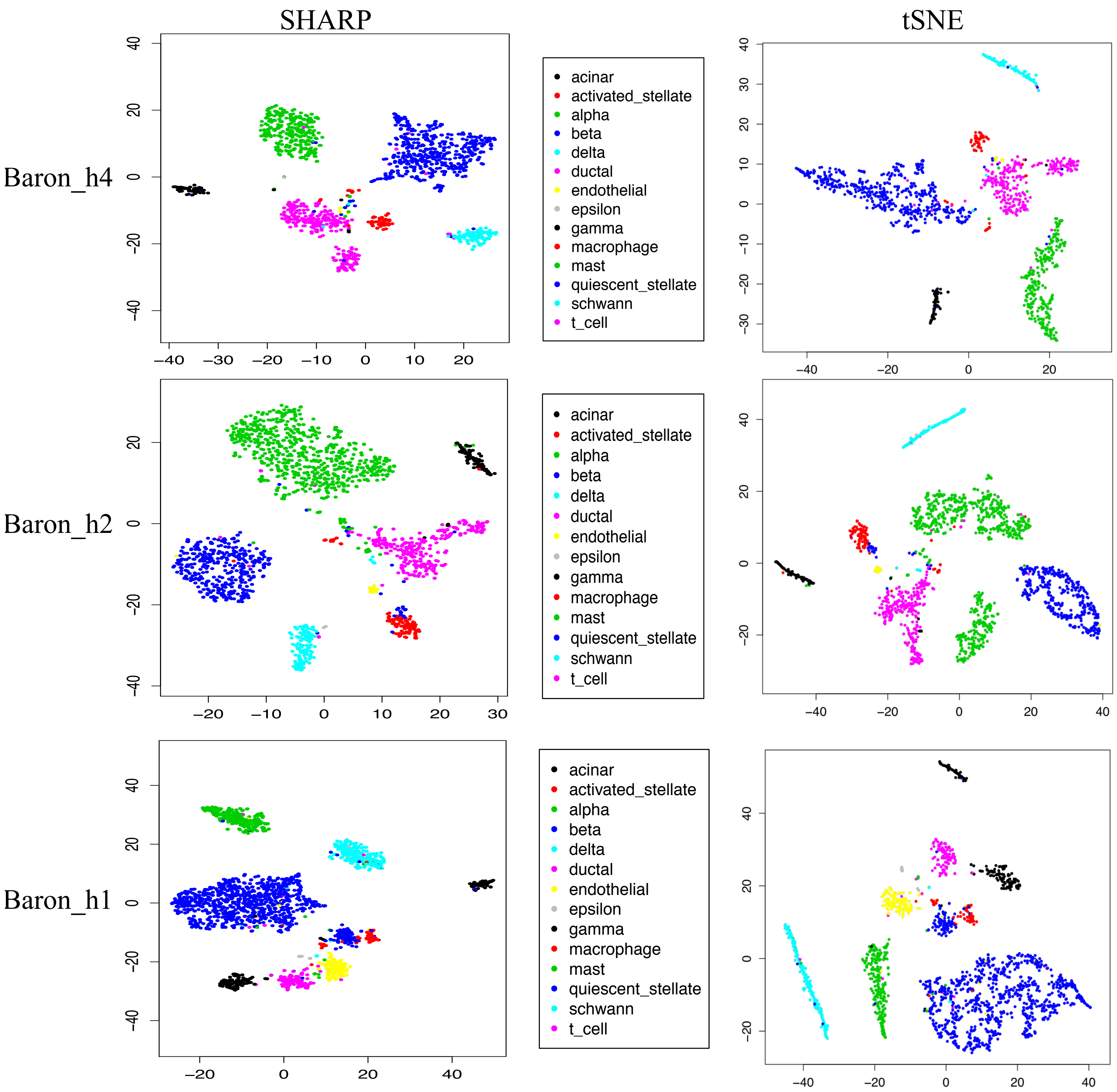


**Supplemental Fig S6c**: Visualization capabilities of SHARP compared with that of tSNE in the Baron_h4 [18] and Baron_h2 [18] and Baron_h1 [18] datasets. The numbers of single cells in these three datasets are 1303, 1724 and 1937, respectively.

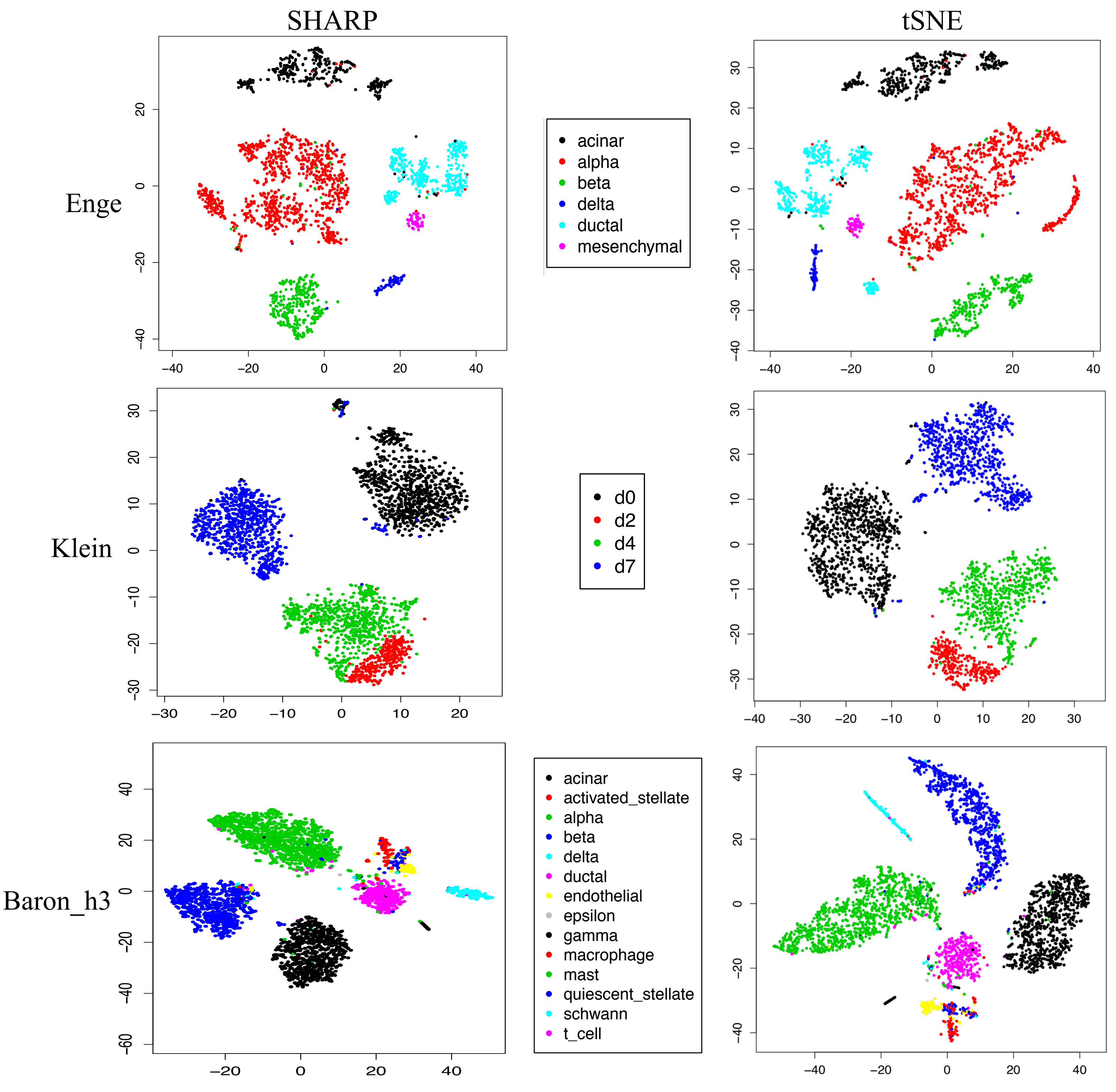


**Supplemental Fig S6d**: Visualization capabilities of SHARP compared with that of tSNE in the Enge [19], Klein [20] and Baron_h3 [18] datasets. The numbers of single cells in these three datasets are 2282, 2717 and 3605, respectively.
