## Supplemental Fig S7 for "SHARP: Single-cell RNA-seq Hyper-fast and Accurate Processing via Ensemble Random Projection"

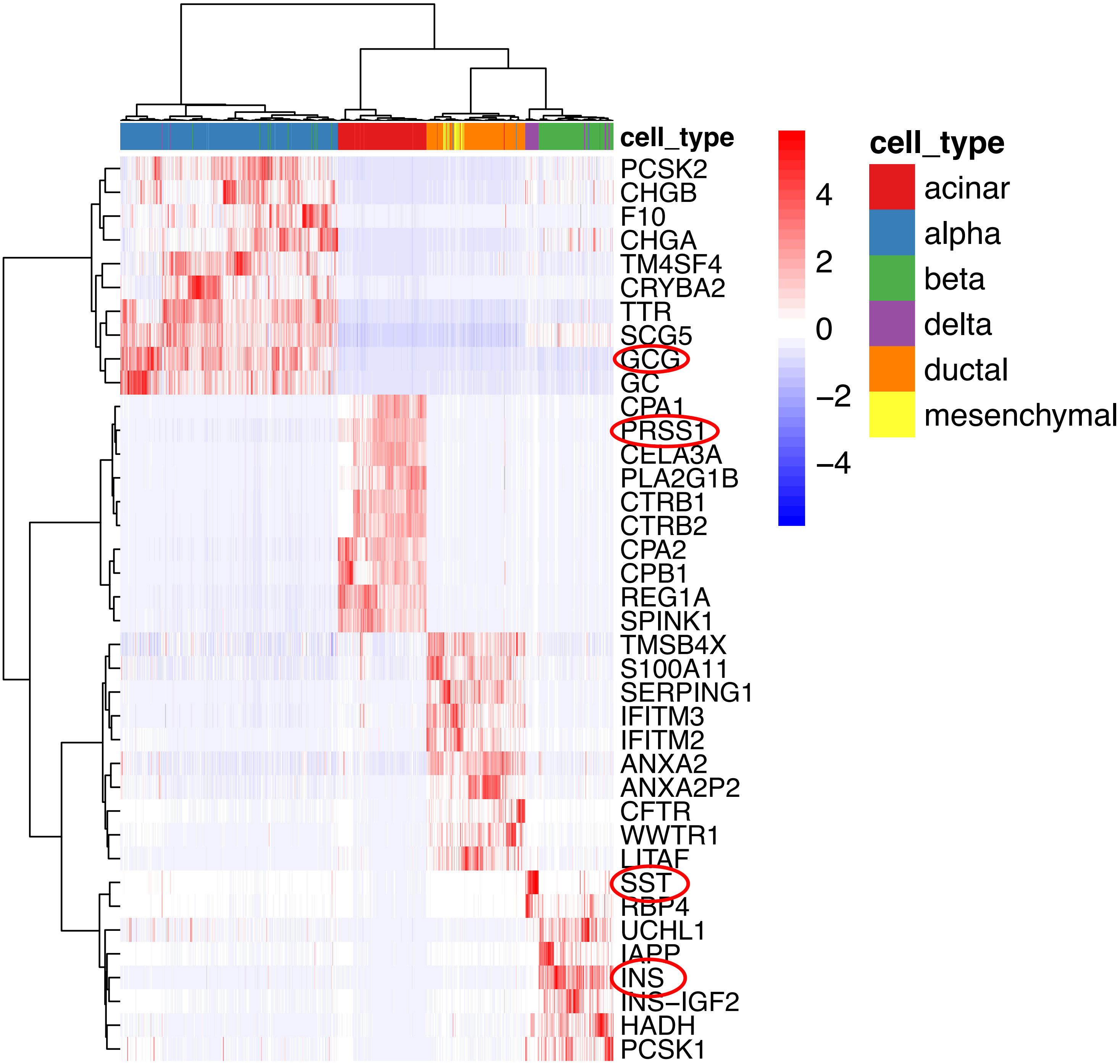


**Supplemental Fig S7**: Cluster-specific marker genes detected by SHARP for the Enge et al. dataset [19]. Each row and each column of the heat-map represent a gene and a cell, respectively. Cells are marked by their original cell types: acinar, alpha, beta, delta, ductal and mesenchymal cells.
