## Supplemental Fig S8 for "SHARP: Single-cell RNA-seq Hyper-fast and Accurate Processing via Ensemble Random Projection"

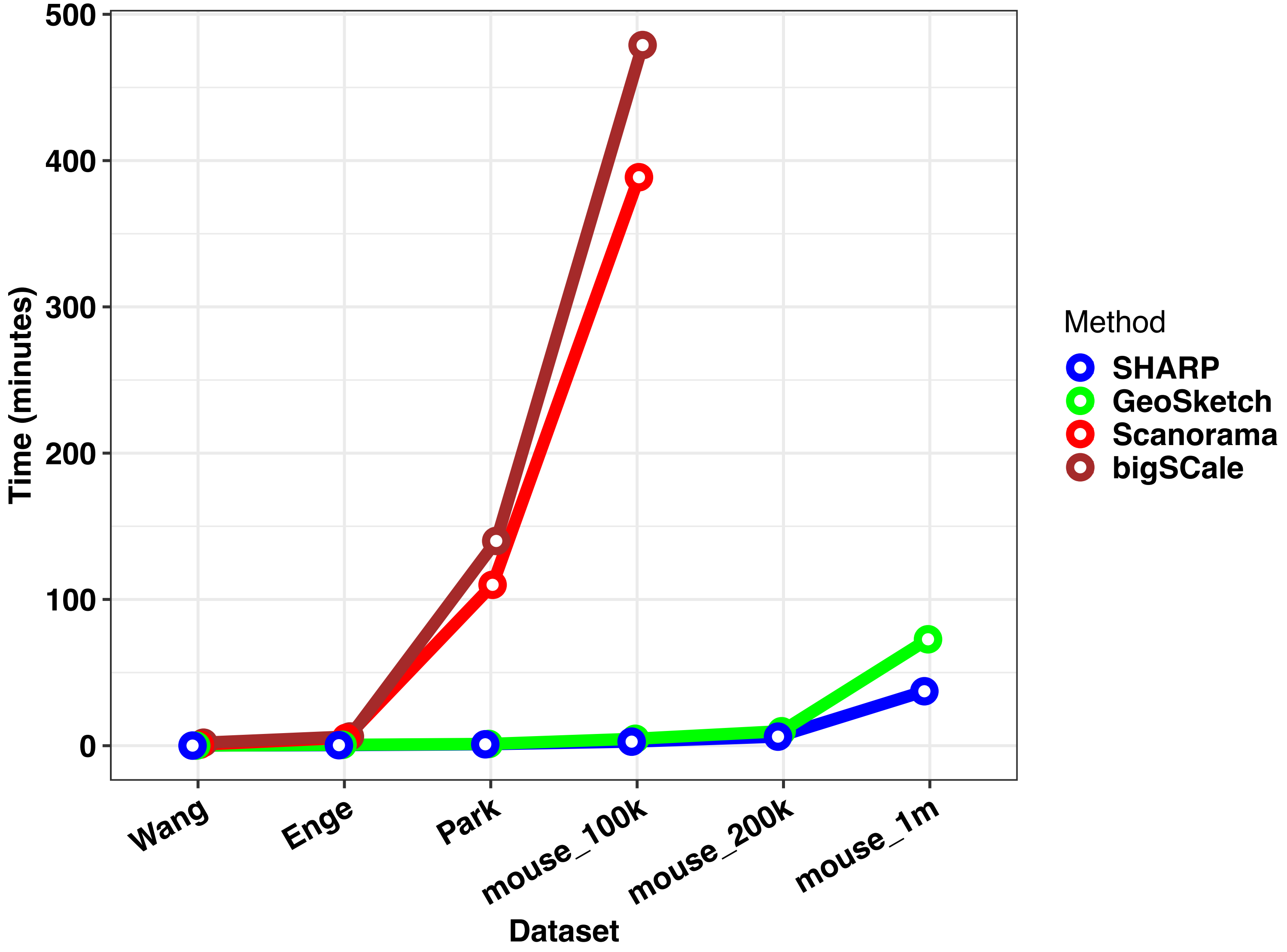


**Supplemental Fig S8**: Comparing SHARP with state-of-the-art scalable scRNA-seq methods in terms of scalability. Geometric Sketching (GeoSketch) is for selecting representative cells from large-scale scRNA-seq datasets, Scanorama is for scRNA-seq data integration and batch correction, and bigSCale is for clustering and marker gene identification. Here, we selected 6 representative scRNA-seq datasets for testing, namely Wang, Enge, Park, mouse_100k, mouse_200k and mouse_1m, with cell numbers of 479, 2282, 43745, 100000, 200000, 1000000, respectively. Among them, the last three datasets were generated by randomly sampling from the 1.3 million cells. For Scanorama, we randomly divided the scRNA-seq data of interest into 2 sub-datasets, each with roughly half of the original cell numbers, and then tried Scanorama on them.
