## Supplemental Fig S10 for "SHARP: Single-cell RNA-seq Hyper-fast and Accurate Processing via Ensemble Random Projection"

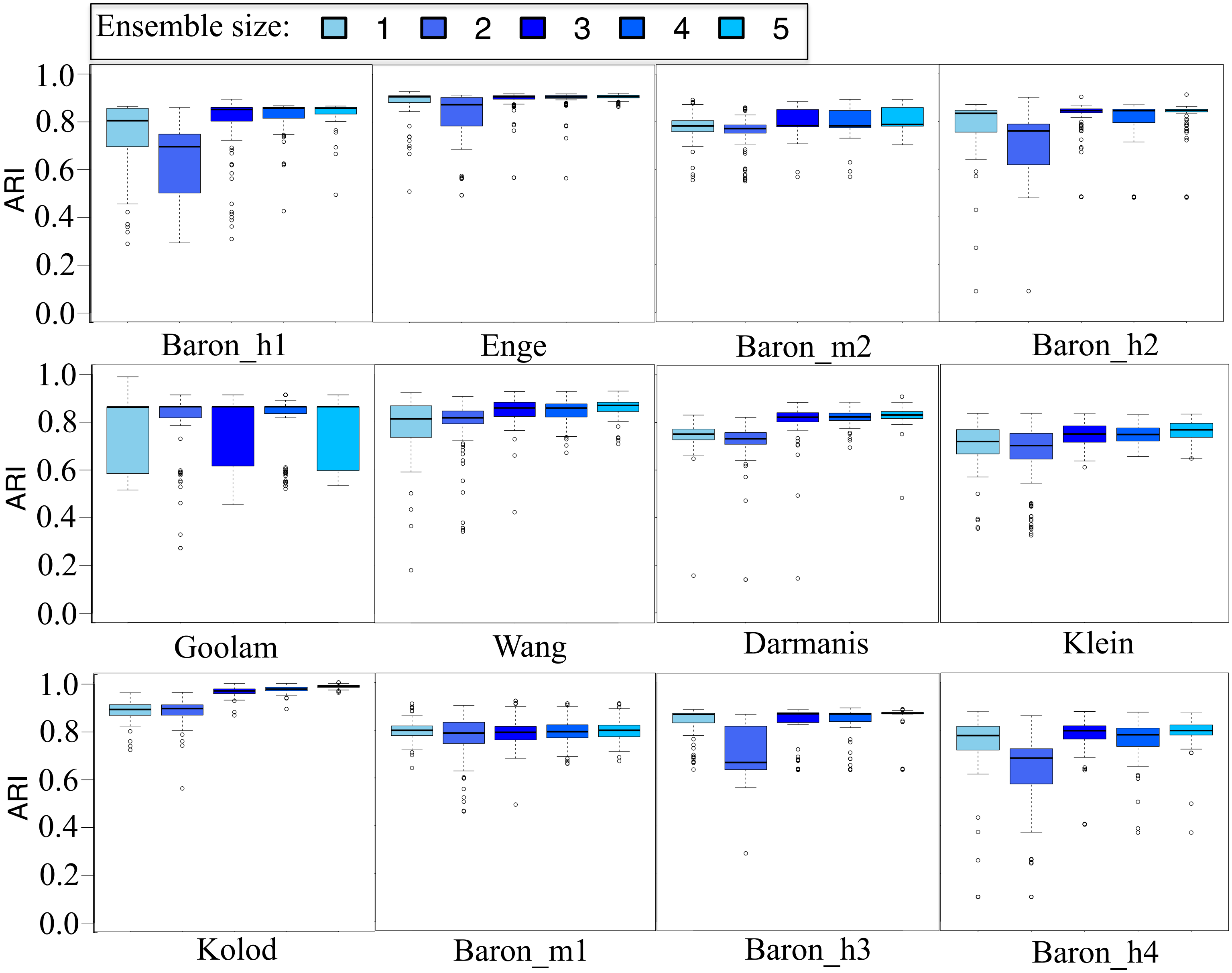


**Supplemental Fig S10**: Performance of SHARP using different ensemble sizes ranging from 1 (i.e., individual RP) to 5 on the 12 datasets, including Baron_h1 [18], Enge [19], Baron_m2 [18], Baron_h2 [18], Goolam [14], Wang [16], Darmanis [15], Klein [20], Kolod [17], Baron_m1 [18], Baron_h3 [18] and Baron_h4 [18]. All of the results are based on 100 runs of SHARP on each corresponding dataset.
