## Supplemental Fig S12 for "SHARP: Single-cell RNA-seq Hyper-fast and Accurate Processing via Ensemble Random Projection"

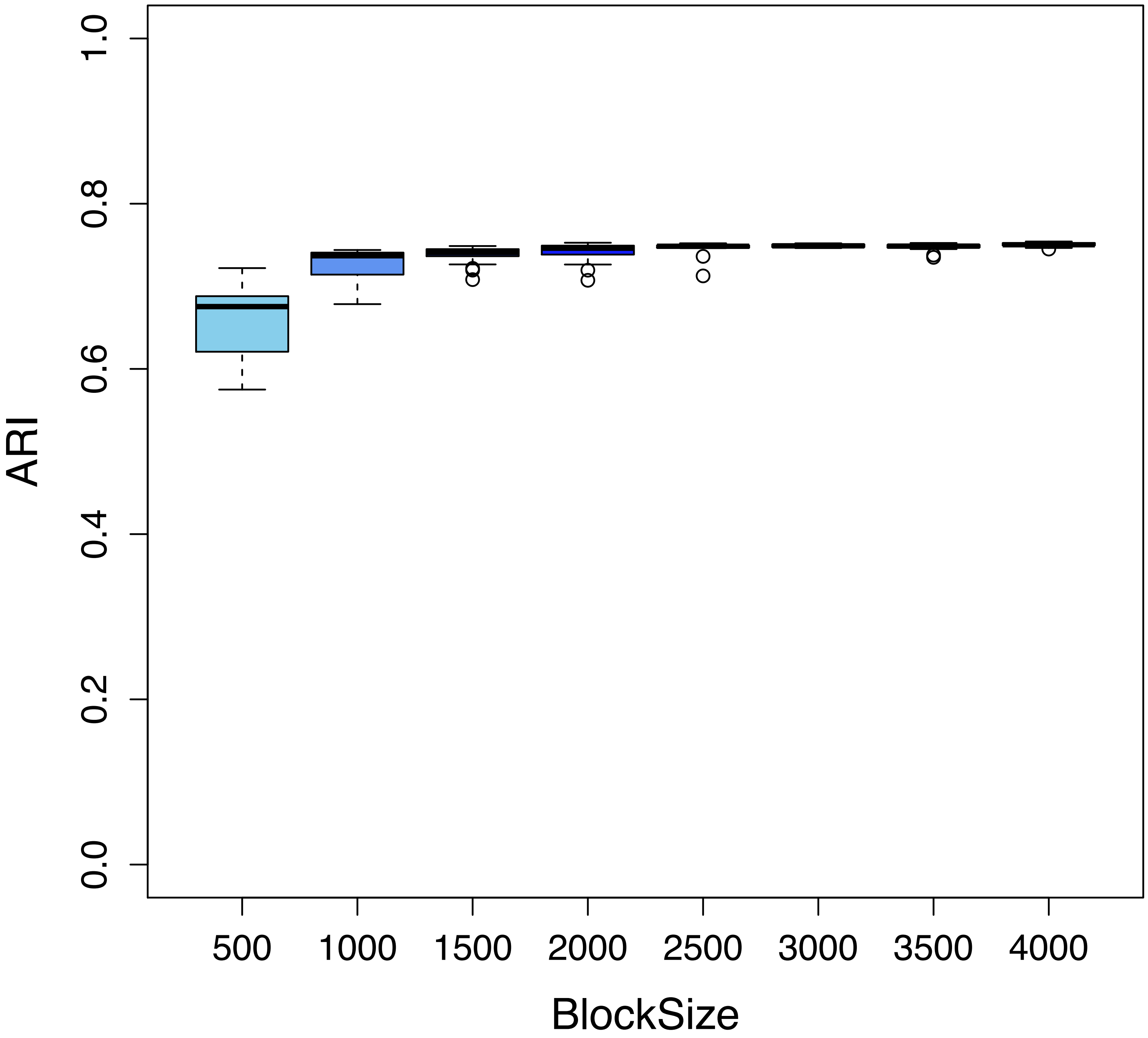


**Supplemental Fig S12**: Performance of SHARP using different block sizes on Park et al dataset [18].BlockSize: number of single cells in each block. The results are based on 100 runs of SHARP on the dataset.
