## Supplemental Fig S13 for "SHARP: Single-cell RNA-seq Hyper-fast and Accurate Processing via Ensemble Random Projection"

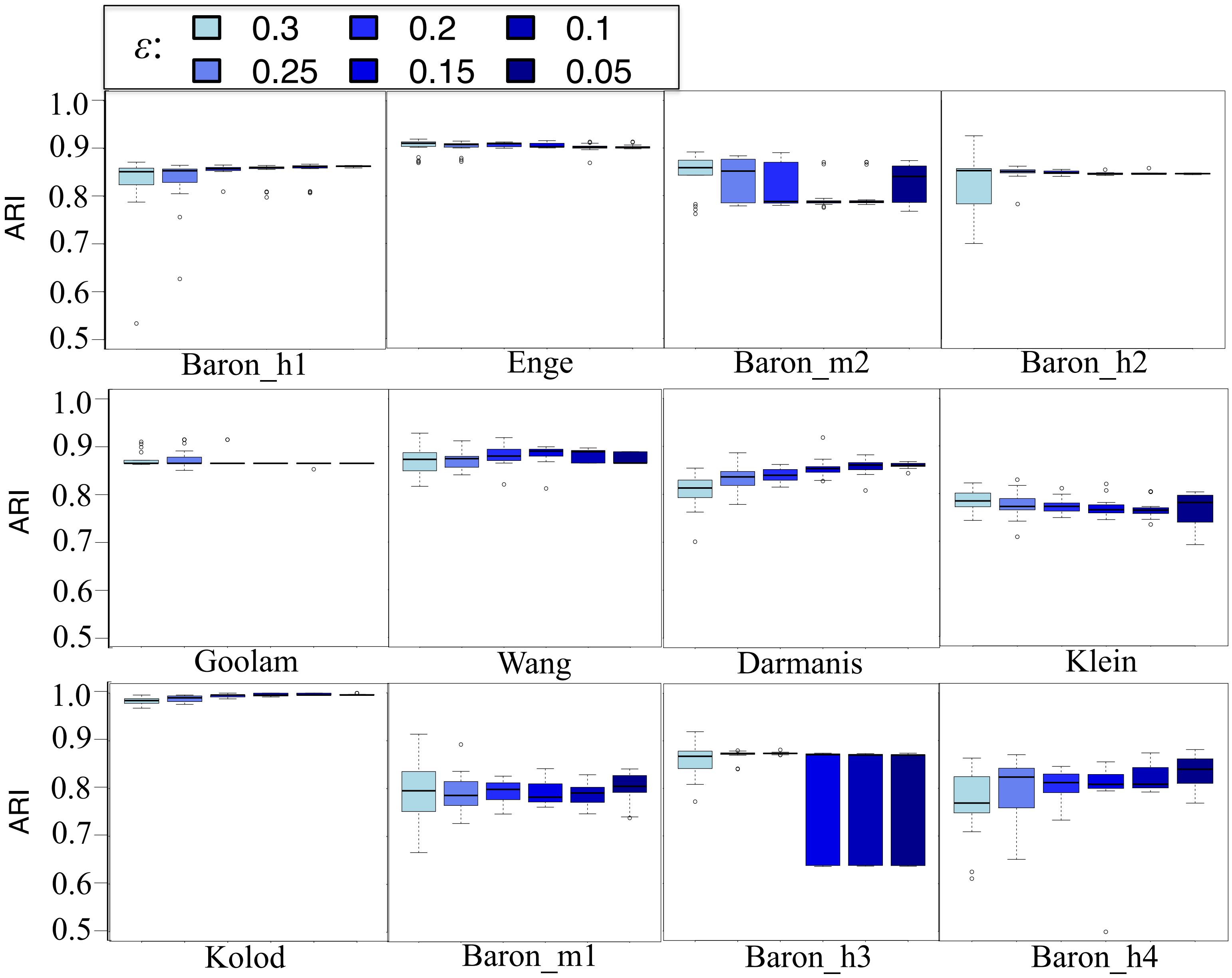


**Supplemental Fig S13**: Comparing SHARP using different values of epsilon (or reduced dimensions) in terms of clustering performance on 12 datasets. All of the results are based on 100 runs of SHARP on each corresponding dataset.
