## Supplemental Fig S14 for "SHARP: Single-cell RNA-seq Hyper-fast and Accurate Processing via Ensemble Random Projection"

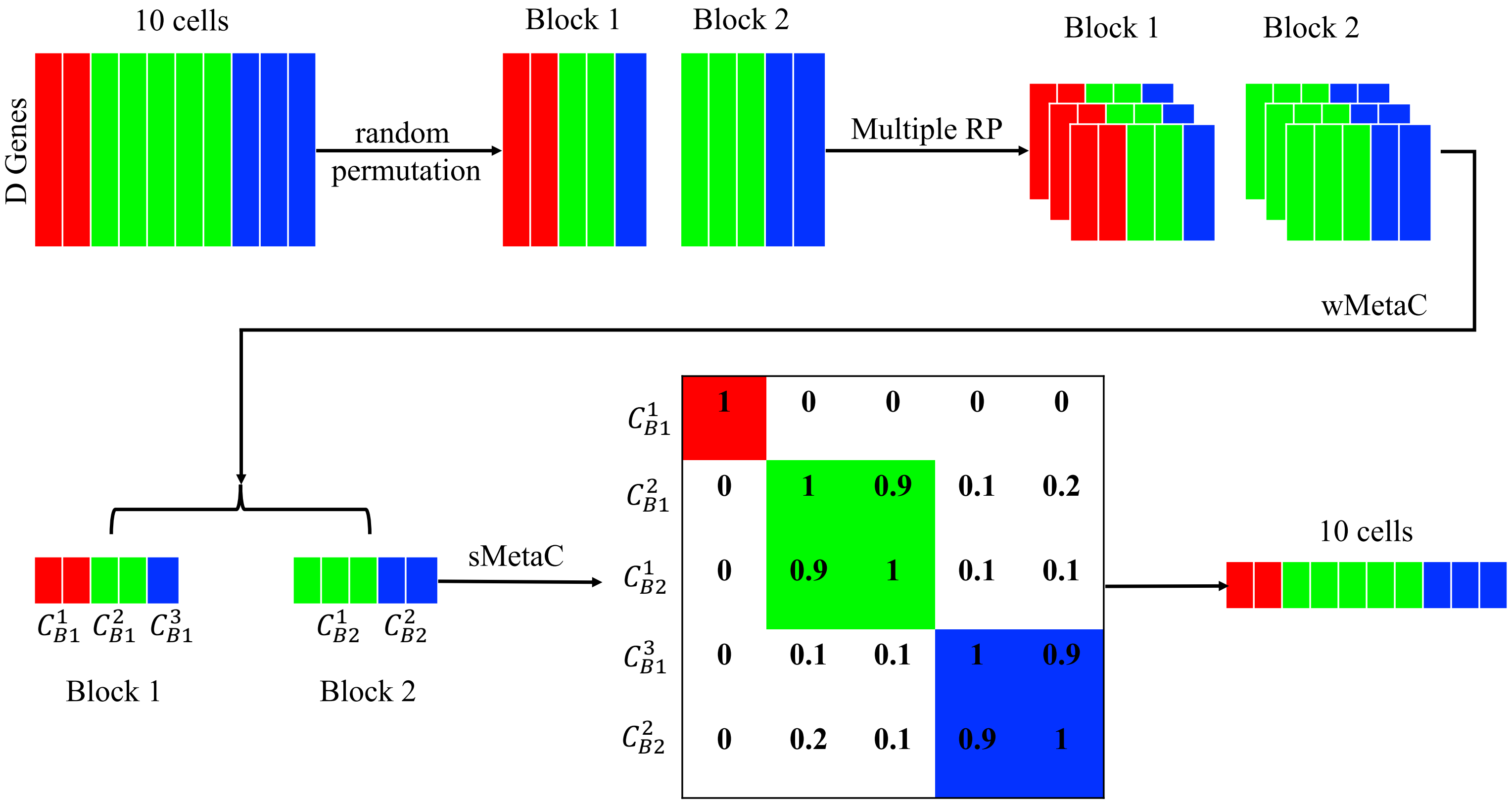


**Supplemental Fig S14**: An example showing how SHARP identify and handle tiny clusters for large-scale scRNA-seq datasets. Suppose the original scRNA-seq dataset consists of 10 cells, which are distributed in three clusters, 2 in Red, 5 in Green and 3 in Blue. It goes through the following stages: random permutation, multiple runs of RP, wMetaC and sMetaC. After random permutation, there is one tiny cluster (with only one cell), namely the Blue cluster in Block 1.
