## Supplemental Fig S15 for "SHARP: Single-cell RNA-seq Hyper-fast and Accurate Processing via Ensemble Random Projection"

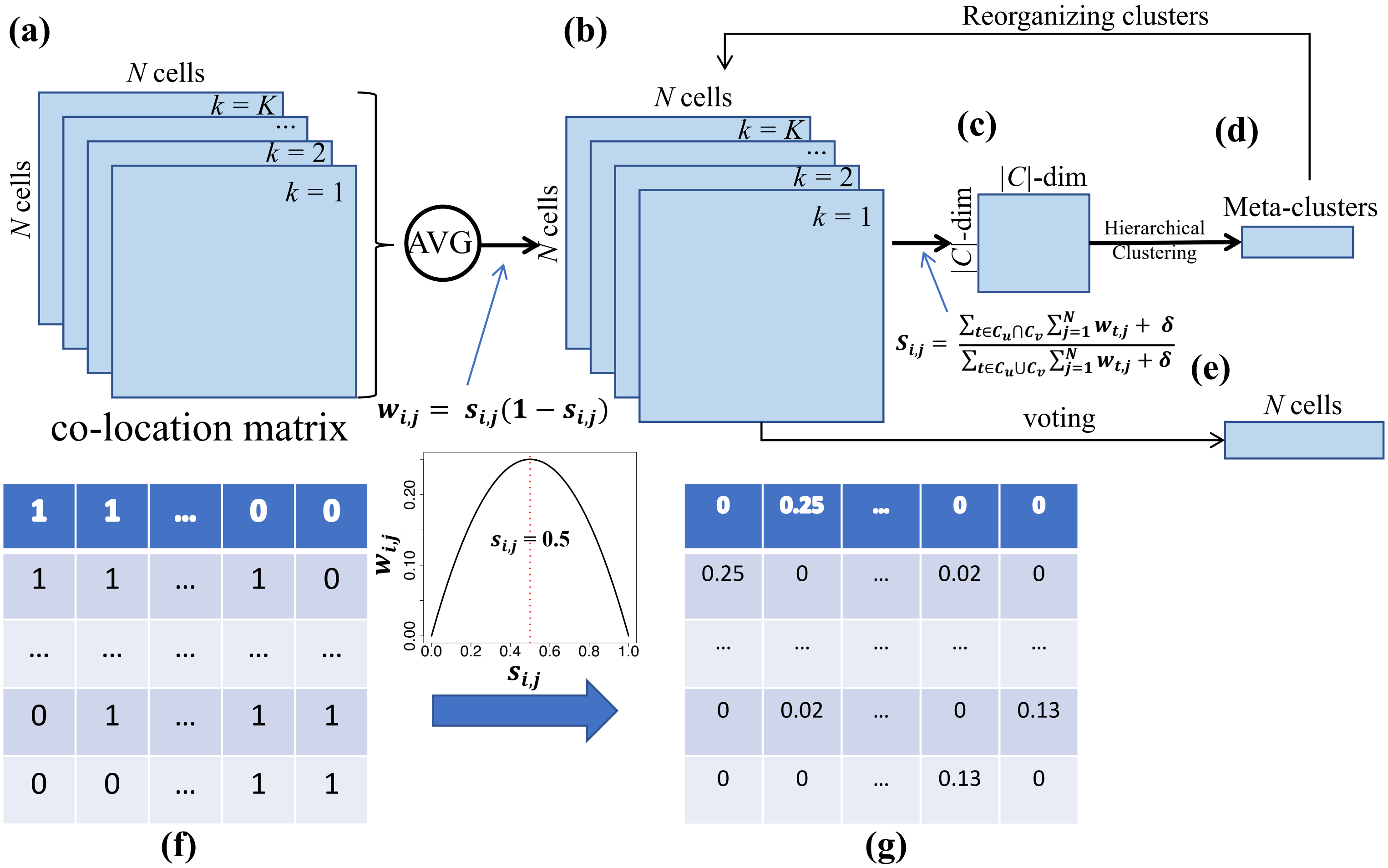


**Supplemental Fig S15**: Flowchart of wMetaC ensemble clustering method. (a) cell-to-cell matrices of the individual clustering results; (b) the weighted cell-to-cell matrices of the individual clustering results; (c) the meta-clustering matrix whose element represents the similarity between each cluster from the weighted individual clustering results; (d) the meta-clustering results by a hierarchical clustering; (e) the final clustering results after voting based on the reorganized individual clustering results; (f) an example of the original cell-to-cell co-location matrix of the individual clustering results, in which 1 indicates that the corresponding two cells are located in the same cluster, whereas 0 means that they are not in the same cluster; (g) an example of the weighted cell-to-cell co-location matrix of individual clustering results where the higher the value of each element, the more difficult to cluster the corresponding two cells. See Online Methods for more detailed descriptions.
