## Supplemental Methods for "SHARP: Single-cell RNA-seq Hyper-fast and Accurate Processing via Ensemble Random Projection"

### Supplemental Note 1: Details of comparing SHARP with state-of-the-art methods methods

SIMLR is a kernel-based machine learning method for clustering scRNA-seq data. It learns cell-to-cell similarity by optimizing a multi-Gaussian-kernel based objective function. SIMLR is based on strong mathematical reasoning which approximates to the theoretical optimized solutions and it also considers regularization to avoid over-fitting. However, because SIMLR needs to calculate cell-to-cell similarity, it is not applicable or at least computational intractable when the number of single cells is very large. Also, the optimization solution is an iterative process which requires considerable time to process.

Compared to SIMLR, SHARP divides the large-scale datasets into blocks of smaller-size single cells. Also, SHARP uses RP to reduce the dimensionality to further expedite the clustering procedures without compromising the clustering performance.

SC3 is a k-means based ensemble clustering method. The general procedures for SC3 include the following steps: (1) gene filtering; (2) PCA or Laplacian transformation for dimension reduction; (3) several runs of k-means clustering and then (4) consensus (or ensemble) clustering by the cluster-based similarity partitioning algorithm (CSPA). The disadvantages of SC3 is that k-means clustering may yield significantly unstable results due to different initializations. Therefore, to achieve relatively robust performance, SC3 requires a significantly large number of iterations (i.e., $10^9$ iterations) for k-means at the expense of long-time processing.

Compared to SC3, SHARP has the following advantages. First, for dimension reduction, SHARP uses random projection, which has a better cell-to-cell distance-preserving property than PCA in the projected low-dimensional space. This can guarantee that the former produces more robust performance than the latter. Second, RP-based clustering runs faster than PCA-based SC3 (Fig. 1b). (3) the weighted meta-clustering (wMetaC) used in SHARP is more powerful than CSPA in terms of ensemble clustering because the latter treats all instances (single cells) with the same weights whereas the former assigns larger weights to those difficult-to-cluster single cells so that performance can be improved (Fig. 1c).

Hierarchical clustering (hclust) is a standard clustering method. hclust is deterministic. The clustering results are organized in a hierarchical tree which can be easily clustered into a designated number of clusters. However, given a large-scale high-dimensional scRNA-seq data, hclust is usually time-consuming and sometimes can come across memory overflow problems. Moreover, hclust is also sensitive to outliers or noise. Besides, sometimes it is not easy to determine the number of clusters from the dendrogram.

Compared to hclust, SHARP has successfully addressed the aforementioned problems. First, SHARP uses a divide-and conquer strategy to partition the large-scale scRNA-seq data into several blocks of small-size data and then utilizes RP to remarkably reduce the dimensions for each block, which makes the processing and analysis of scRNA-seq data much faster and simultaneously avoids memory overflow problems. Second, by using sparse RPs, many of the outliers or noises are smoothed out, which are good for clustering. Third, the number of clusters is determined by integrating three different indices (i.e., Silhouette index, CH index and hierarchical heights).

tSNE is a nonlinear dimensionality reduction technique which projects scRNA-seq data with thousands or tens of thousands of dimensions into a 2-D or 3-D space. In tSNE space, similar instances (i.e., single cells) are modeled by nearby points and dissimilar instances are modeled by distance points. A popular clustering method is to use k-means to cluster the 2-D or 3-D tSNE-based feature matrix. tSNE are (1) fast, (2) simple and straightforward, and (3) easy to implement. However, disadvantages are that tSNE can only preserve the nearest neighbors, but can not preserve cell-to-cell distances nor density. Besides, k-means based clustering will yield unstable performance.

Compare to tSNE plus k-means, SHARP uses RP for dimension reduction, which can well preserve the cell-to-cell distances in the low-dimensional space. By using ensemble RP, SHARP can produce more robust and better clustering performance. Meanwhile, it is noted that practically PCA is usually done before tSNE (e.g., the Rtsne package will use PCA first to reduce the original dimensions to a smaller dimension, for example, 50, and then tSNE is used for nonlinear dimension reduction). Therefore, SHARP runs faster than tSNE plus k-means.

Similar to tSNE, UMAP is also a non-linear dimension reduction technique. The difference between tSNE and UMAP is that the latter uses a manifold learning method which is based on Riemannian geometry and algebraic topology. UMAP is competitive with tSNE in terms of visualization and is alleged to preserve more of the global structure which is faster than tSNE.

CIDR, which is short for clustering through imputation and dimensionality reduction, is an algorithm which uses an implicit imputation approach to alleviate the impact of dropouts in scRNA-seq data. Essentially speaking, CIDR is an improved PC-like algorithm which is faster than PCA and more suitable for dealing with the dropouts in scRNA-seq data. However, our experimental results suggest that CIDR is only fast when the number of cells is relatively small (e.g., < 10000). When the number of cells is large (e.g., > 10000), the running time for CIDR is exponentially increased.

Phenograph is a high-dimensional data clustering method which works by creating a graph representing cell-to-cell similarities and then identifying communities in the graph. The graph is built in a two steps: using a k-nearest neighbor method to find the k neighboring cells for each cells and then build weighted graphs by scaling the cell-to-cell weight with the number of sharing neighbors. Phenograph is generally not scalable to large-scale scRNA-seq data clustering (See Fig. 1b and Fig. 1c in the main paper) because calculating the distance in the kNN methods will yield significantly increased time complexity when the number of cells is very large. Another disadvantage of this method is that kNN is susceptible to the noisy data and thus Phenograph may not be robust when the noisy data imposes a relatively large impact on the data.

Seurat is an integrated package which is not only used for clustering but also designed for quality control, analysis and exploration of single-cell RNA-seq data. It uses multiple interesting algorithms like canonical correlation analysis and the so-called nonlinear warping algorithm for gene correlation structure learning and feature scaling normalization. It is relatively faster because it just uses the most variant genes, which, however, may ignore important information for clustering. Although Seurat is a multi-functional package, it performs poorly in terms of clustering performance (See Fig. 1c and Supplemental Fig. 3).

### Supplementary Note 2: Adjusted Rand Index (ARI)

The Adjusted Rand index (ARI) is a corrected-for-chance version of the Rand index, a similarity measurement between a predicted clustering and the corresponding reference (e.g., ground-truth or published) clustering. Generally speaking, ARI is to measure how accurately a prediction of clustering is made in the unsupervised learning scenarios, which is similar to the accuracy measurement in supervised classification problems.

In our case, the reference clustering for a given single-cell RNA-seq (scRNA-seq) dataset (e.g., with $N$ single cells) is available (i.e., those clusters or sub-populations identified in the corresponding published datasets). For ease of presentation, we denote the given reference clustering as : $\mathscr{R} = \{\mathscr{R}_1, \ldots, \mathscr{R}_s, \ldots, \mathscr{R}_S\}$, where the set $\mathscr{R}$ has $S$ clusters and $\{\mathscr{R}_s\}_{s=1}^{S}$ represents the set of single cells in the $s$-th cluster. Similarly, for the corresponding predicted clustering by SHARP, we denote it as:$\mathscr{P} = \{\mathscr{P}_1, \ldots, \mathscr{P}_t, \ldots, \mathscr{P}_T\}$, where the set $\mathscr{P}$ has $T$ clusters and $\{\mathscr{P}_t\}_{t=1}^{T}$ represents the set of single cells in the $t$-th cluster. In this case, we further denote the number of single cells shared by the $s$-th cluster $\mathscr{R}_s$ in the reference clustering and the $t$-the cluster $\mathscr{P}_t$ in the predicted clustering as: $c_{s,t} = |\mathscr{R}_s \cap \mathscr{P}_t|$, where $|\cdot|$ and $\cap$ denote the set cardinality and set intersection, respectively. Thus, the ARI can be formulated as follows:

$$
\begin{aligned}
\text{ARI} &= \frac{\text{Rand Idex - Expected Index}}{\text{Max Index - Expected Index}} \\
&= \frac{\sum_{s=1}^{S}\sum_{t=1}^{T}\binom{c_{s,t}}{2} - \left[\sum_{s=1}^{S}\binom{x_s}{2}\sum_{t=1}^{T}\binom{y_t}{2}\right]\Big/\binom{N}{2}}{\frac{1}{2}\left[\sum_{s=1}^{S}\binom{x_s}{2} + \sum_{t=1}^{T}\binom{y_t}{2}\right] - \left[\sum_{s=1}^{S}\binom{x_s}{2}\sum_{t=1}^{T}\binom{y_t}{2}\right]\Big/\binom{N}{2}}\,,
\end{aligned}
\tag{S1}
$$

where $x_s = \sum_{t=1}^{T} c_{s,t}$ , $y_t = \sum_{s=1}^{S} c_{s,t}$ and $\binom{k}{2} = \frac{k(k-1)}{2}$ is a binomial coefficient.

From Eq. (S1), we can see that compared to RI, ARI reduces the random chance (i.e., expected index) of high-degree agreement between the reference clustering and the predicted clustering from RI, thus making the former much more reliable than the latter.

It should be noted that the value of ARI can be negative and the maximum value is +1. Generally, the larger ARI, the better the predicted clustering, with +1 indicating that the predicted clustering is perfectly consistent with the reference clustering, whereas 0 (negative) indicating that the predicted clustering is as good as random guess (worse than random guess).

### Supplemental Note 3: Single-core vs multi-core

To make full use of computational resources, SHARP is implemented in a parallelization way, i.e., using multi cores to accelerate the processing of scRNA-seq data. We compared the performance of SHARP over SC3 and SIMILR on the Montoro et al. dataset [17] by using single-core and multi-core configurations. By default, the SHARP package will use (*n*-1) cores, where *n* is the number of CPU cores. In Supplemental Figure 1, we used 16 cores for all of the three algorithms. All of results for SHARP and SC3 are based on 100 runs of the corresponding algorithms, because these algorithms are stochastic. For SIMLR, the results are deterministic and thus one run is sufficient. Note that due to the large size of the Montoro dataset (i.e., 66,000+), the original version of SIMLR and SC3 has memory over-flow problems, thus not capable of clustering. Thus, SIMLR was equipped with k-nearest-neighbor algorithm, whereas SC3 with support vector machines (SVM).

### Supplemental Note 4: Cell-to-cell distance

To investigate the distance-preserving properties of SHARP in low-dimensional space, we compared the cell-to-cell distance SHARP with PCA and tSNE for 12 single-cell datasets (Supplemental Figure 2). Specifically, we first calculated the cell-to-cell distances using the pairwise Pearson correlations between each pair of cells for the original scRNA-seq data, the PCA-projected data, tSNE-projected data and SHARP-projected data. Then, we calculated the correlations between each pair of these four kinds of distance-based data and simultaneously did the correlation tests. Note that the dimension of tSNE-projected data is 3 because tSNE generated 2-D or 3-D data for visualization (we selected the 3-D). For SHARP, the projected data is the ensemble of each RP-projected data, where the ensemble size is 5. For fair comparison, the reduced dimension for PCA is the same as that used for SHARP. All of the correlation coefficients here are the Pearson correlation coefficients.

### Supplemental Note 5: Ensemble RP vs individual RP

One run of RP may yield diverse and volatile performance. Therefore, ensemble of RP-based clustering is necessary for achieving robust performance. We compared ensemble RP (SHARP) with individual RP on 12 datasets (Supplemental Figure 4). By default, for small-size datasets (e.g., < 5,000 single cells), the ensemble RP is based on 15 applications of individual RPs, whereas for large-size datasets (e.g., >5,000 single cells), we used 5 applications of RPs. The results of ensemble RP are based on 100 runs of SHARP on each dataset; in other words, the results of individual RP are based on 1,500 runs of RP-based clustering. It should also be noted that the ensemble part is to combine the results of using hierarchical clustering (hclust) on the RP-based dimension-reduced feature matrix for each run, rather than to simply make ensemble of the RP-based feature matrices and then use hclust on the ensemble matrix. This is because similar to classification problems (Wan et al. 2014), scoring fusion (the former) generally produces better performance than feature fusion (the latter).

### Supplemental Note 6: Ensemble size, block size and subspace dimensions

#### Ensemble size

SHARP combines the results of several runs of individual RPs, thus the ensemble size (i.e., the number of runs of RPs) is a critical parameter in the performance of SHARP. To investigate the effect of ensemble size on SHARP, we have tried ensemble sizes from 5 to 50 for 12 single-cell datasets (Supplemental Figure 5). In our R package of SHARP, the default ensemble size for small-size single-cell datasets is 15, whereas the parameter for large-scale datasets (larger than 5,000) is 5.

#### Block size

When the number of single cells is large (e.g., > 5,000), data partition, or partitioning scRNA-seq data into several blocks, is necessary to guarantee fast processing of scRNA-seq data by SHARP. The block size is defined as the base number of single cells in each block, i.e., a user-defined parameter (the parameter *n* in Section "Data partition" of Online Methods) which is the upper bound of each block of single cells. To investigate how the block size affects the performance of SHARP, we implemented SHARP using different block sizes ranging from 500 to 4,000 on the Park dataset with 43,000+ single cells (Supplemental Figure 6). Too many clusters as input to the ensemble clustering will consume much more time which overshadows the advantages of using small block size. On the other hand, the clustering performance of SHARP also gets improved when the block sizes increases. Therefore, a proper block size should be selected to strike a balance between fast processing and robust clustering performance.

#### Subspace dimension

RP is essentially a dimension-reduction technique. Thus, how much the dimensionality should be reduced is a key question when applying it to single cell analysis. The subspace dimension or the reduced dimensionality $d$ in SHARP is controlled by a parameter $\epsilon$ in the equation $d = \log_2(N)/\epsilon^2$. To investigate how the reduced dimensions affect the performance of SHARP, we have tried different $\epsilon$ (from 0.05 to 0.3) for 12 single-cell datasets. The larger $\epsilon$, the smaller the subspace dimension. We implemented SHARP with different $\epsilon$ (i.e., different reduced dimensions) on 12 datasets (Supplemental Figure 7).

### Supplemental Note 7: How SHARP handles tiny clusters

To elaborate how SHARP handles tiny clusters generated by the random permutation step, Supplemental Fig S14 was given to exemplify the process. Specifically, suppose the original scRNA-seq consists of 10 cells, which are distributed in three clusters, 2 in Red, 5 in Green and 3 in Blue. After random permutation, the 10 cells are divided into 2 blocks, i.e., Block 1 (2 in Red, 2 in Green and 1 in Blue) and Block 2 (3 in Green and 2 in Blue). So the Blue cluster in Block 1 is an example of a tiny cluster (with only one cell). Then, multiple random projections are performed for each block followed by the wMetaC approach so that each block is clustered. After wMetaC, Block 1 is grouped into 3 clusters, i.e., $C_{B1}^{1}$ (2 cells), $C_{B1}^{2}$ (2 cells) and $C_{B1}^{3}$ (1 cell), and Block 2 is grouped into two clusters, i.e., $C_{B2}^{1}$ (3 cells) and $C_{B2}^{2}$ (2 cells). Then in the sMetaC step, the pairwise similarities among these 5 clusters were calculated. Based on the pairwise similarities, the 5 groups were meta-clustered into 3 clusters, $\{C_{B1}^{1}\}$, $\{C_{B1}^{2}, C_{B2}^{1}\}$ and $\{C_{B1}^{3}, C_{B2}^{2}\}$. When mapping back, the original 10 cells were grouped into 3 clusters, 2 in Cluster 1 (i.e., Red), 5 in Cluster 2 (i.e., Green) and 3 in Cluster 3 (i.e., Blue). In this case, the tiny clusters were identified, and then were merged based on the meta-cluster-similarity in the sMetaC step.

### Supplemental Note 8: Johnson-Lindenstrauss Lemma

Random projection (RP) is a group of simple yet powerful dimension-reduction technique. It is based on the Johnson-Lindenstrauss lemma (Johnson and Lindenstrauss 1984) given below:

**Lemma 1.** Given $\epsilon > 0$, a set $\mathscr{X}$ of $N$ points in $\mathscr{R}^T$, and a positive integer $d \geq d_0 = \mathscr{O}(\log N/\epsilon^2)$, there exists such that

$$(1-\epsilon)||\mathbf{u}-\mathbf{v}||^2 \leq ||\mathbf{f}(\mathbf{u})-\mathbf{f}(\mathbf{v})||^2 \leq (\mathbf{1}+\epsilon)||\mathbf{u}-\mathbf{v}||^2$$

for all $\mathbf{u}, \mathbf{v} \in \mathscr{X}$.

The lemma suggests that if points in a high-dimensional space are projected onto a randomly selected subspace of suitable dimension, the distances between the points are approximately preserved (Frankl and Maehara 1988).

Specifically, the original *D*-dimensional data is projected onto a *d*-dimensional subspace, using a random matrix whose column are unit length, i.e.,

$$\mathbf{P} = \frac{\mathbf{1}}{\sqrt{\mathbf{d}}}\mathbf{RM} \in \mathscr{R}^{d\times N}, \mathbf{M} \in \mathscr{R}^{D\times N}, \mathbf{R} \in \mathscr{R}^{d\times D}$$

As long as the elements of **R** conforms to any distributions with zero mean and unit variance, **R** gives a mapping that satisfies the Johnson-Lindenstrauss lemma.

### Supplemental Note 9: Determining number of clusters

SHARP determines the number of predicted clusters by using three criteria: Silhouette index (Rousseeuw 1987), Calinski-Harabasz (CH) index (Caliński and Harabasz 1974) and hierarchical heights. When the Silhouette index is with high score, we determine the number of clusters as the one with the maximum average Silhouette index; if the Silhouette index is not with high score, we resort to the CH index to determine the cluster number; if both indices are not with high confidence, we use the hierarchical heights to determine the predicted number of clusters. Now we go into details about each of them.

#### (a) Silhouette Index

Unlike adjusted Rand index (ARI) which is an external measurement based on gold standard (or reference) clustering to determine how good a clustering prediction is, the Silhouette index is an internal measurement which is evaluated solely based on the clustering itself. Therefore, it is a potentially good index to automatically determine the optimal number of clusters for a clustering method. Conceptually speaking, the Silhouette index contrasts the average within-cluster distance with the average between-cluster distance, thus the larger the Silhouette index, the better the clustering performance.

Specifically, for a predicted clustering by SHARP, e.g., $\mathscr{P} = \{\mathscr{P}_1, \ldots, \mathscr{P}_t, \ldots, \mathscr{P}_T\}$, suppose the $i$-th single cell $\{s_i\}_{i=1}^{N}$ in the original scRNA-seq data is predicted in the $t$-th cluster $\mathscr{P}_t$. Then, we denote $d_i^{\text{wc}}$ as the average distance between $s_i$ and the other single cells within the same cluster $\mathscr{P}_t$, and $d_i^{\text{bc}}$ as the minimum average distance between $s_i$ and the other single cells in other clusters except $\mathscr{P}_t$. Therefore, the Silhouette Index for the $i$-th single cell $s_i$ is defined as:

$$\begin{aligned}\text{sind}(s_i) &= \frac{d_i^{\text{bc}} - d_i^{\text{wc}}}{\max\left\{d_i^{\text{bc}}, d_i^{\text{wc}}\right\}} \\ &= \begin{cases} d_i^{\text{bc}}/d_i^{\text{wc}} - 1, & \text{if } d_i^{\text{wc}} > d_i^{\text{bc}} \\ 0, & \text{if } d_i^{\text{wc}} = d_i^{\text{bc}} \\ 1 - d_i^{\text{wc}}/d_i^{\text{bc}}, & \text{if } d_i^{\text{wc}} < d_i^{\text{bc}} \end{cases}\end{aligned} \tag{1}$$

From Eq. (1), we can see that $\text{sind}(s_i) \in [-1, +1]$. Obviously, when $\text{sind}(s_i)$ is approaching +1, then $d_i^{\text{wc}}$ is significantly less than $d_i^{\text{bc}}$; in other words, the within-cluster distance is significantly small and the between-cluster distance is significantly large. Thus, a Silhouette index close to +1 means $s_i$ is appropriately clustered; and vice versa. Note that for completeness, for a cluster with only one single cell, the Silhouette index is set to 0.

Usually, the average Silhouette index, namely $\frac{1}{N}\sum_{i=1}^{N} \text{sind}(s_i)$, is used as a measure to determine the optimal number of clusters because it can evaluate how well (or appropriately) the data in whole are clustered. A common way is to select the optimal number of clusters as the one which maximizes the average Silhouette index.

Another reason why the Silhouette index is popularly used is that compared to other internal clustering indices, this index is bounded within [-1, +1], which provides sort of a piece of evidence of how confidently we should trust the Silhouette index. For example, in our case, we choose to believe the selected number of clusters by the Silhouette index when it is larger than 0.35 (by default). Otherwise, we choose another index (the Calinski-Harabasz index discussed in the next Supplemental note) to determine the optimal number of clusters.

The third reason we used the Silhouette index in SHARP is that the indices for different numbers of clusters (e.g., from 2 to 40) can be calculated very fast in the hierarchical clustering (hclust) scenario. This is because after hclust, a tree is produced and we just need to cut the tree into the desired number of groups (or clusters). We can calculate the Silhouette index for each candidate number of clusters (e.g., from 2 to 40) with just running hclust once. Then, based on the obtained Silhouette indices for different numbers of clusters, we select the optimal number of clusters as the one which gives the maximum Silhouette index.

#### (b) Calinski-Harabasz index

When the Silhouette index is smaller than a threshold (e.g., 0.35 by default), it is risky to believe the number of clusters selected by the Silhouette index. In this case, we rely on another internal clustering measurement, namely the Calinski-Harabasz (CH) index. The CH index, also known as the variance ratio criterion, is the weighted ratio between the between-cluster variance and the within-cluster variance. The idea behind the CH index is to simultaneously minimize the within-cluster variance and maximize the between-cluster variance. Compared to the Silhouette index which can be calculated for each single cell, the CH index is an overall index for a whole dataset for different numbers of clusters.

Mathematically, the CH index is defined as:

$$\mathrm{CH} = \frac{1/(C-1)}{1/(N-C)} \cdot \frac{V_{\mathrm{bc}}}{V_{\mathrm{wc}}}, \tag{2}$$

where N is the number of single cells, C is the number of clusters, $V_{\mathrm{bc}}$ and $V_{\mathrm{wc}}$ are the overall between-cluster variance and overall within-cluster variance, respectively. $V_{\mathrm{wc}}$ is similar to the objective function of k-means which measures how tightly the overall clusters are grouped, whereas $V_{\mathrm{bc}}$ evaluate how spreading apart each cluster (especially the centroid of each cluster) is from the other clusters. For the scaling factors, because $V_{\mathrm{bc}}$ is calculated based on the number of clusters which thus has (*C-1*) degrees of freedom, whereas $V_{\mathrm{wc}}$ has (*N-C*) degrees of freedom. To make the $V_{\mathrm{bc}}$ and $V_{\mathrm{wc}}$ on the same scale, we add the proportional scaling factors to them.

Because we want to minimize the within-cluster variance and maximize the between-cluster variance, the larger the CH index, the better the predicted clustering. Unlike the Silhouette index, the CH index is not bounded with a range of values, therefore, it is unlikely to compare the CH index across different datasets. However, we can use the CH index to compare among different number of clusters in the same dataset, thus providing a way to optimize the number of clusters. The largest CH index corresponds to the optimal clustering size.

It should be noted that sometimes the maximum CH index reaches when the cluster size is 2, however, clustering the data into 2 clusters is probably not the ideal case. And a local maximum CH index may correspond to our expected cluster number. In this case, we think the CH index is not reliable and we will resort to another measure, namely the hierarchical heights.

##### (c) Hierarchical heights

When the Silhouette index and the CH index are not working with high confidence, we will use the hierarchical heights to determine the number of clusters. After hierarchical clustering, the data (e.g., single-cell data) are organized into a hierarchical tree or a dendrogram. Any two single cells are eventually linked together at some level in the dendrogram. The height of the link, or the hierarchical height, represents the distance between the two corresponding clusters that contain these two single cells. The hierarchical heights can also be treated as one kind of measurement to how well the dendrogram is constructed by comparing the hierarchical heights with the pairwise distances generated from the original data. If the predicted clustering is good agreement with the reference one, the linking of single cells in the dendrogram should highly correlate with the original distances. Because this is also one of the internal measurements for clustering, it can also be used to determine the number of clusters.

Unlike the previous two indices in which we tried different numbers of clusters to cut the hierarchical tree and then calculated the indices, in this measurement, we use the height to cut the tree and based on the tree-cutting, we can determine the number of clusters. It is reasonable to consider comparing the heights of each link in the hierarchical tree with the heights of neighboring links immediately below it in the tree. Specifically, if the height of a link is approximately the same as that of the link immediately below it, then we can think these links are highly consistent and we should not cut the tree between them. On the contrary, if the height of a link is conspicuously different from that of the links immediately below it (i.e., the height of the current link is significantly larger than that of the immediate next link), then the single cells linked at this level are much farther apart from each other than the other ones. In this case, we should cut the tree between them. We will select the maximum one which satisfies the aforementioned condition.

While in this case, a question arises in terms of where to cut: how different between two consecutive hierarchical heights of the links in the dendrogram should we consider as the criteria to cut the tree? There is no easy answer to this question. However, we do have some principles guiding us to select. First, consider whether we want relatively larger number of clusters or relatively smaller one. Second, make a balance between differentiating the consistent/ inconsistent links and avoiding generating too many clusters by going to too lower levels of trees. Based on these principles, we set the hierarchical height difference between two consecutive links as 2 times, namely, if the current hierarchical height is 2 times larger then the immediate next hierarchical height, we select out those pairs of heights satisfying this condition and then cut the trees at the median of the two maximum heights. For flexibility, we allow users to determine how much (or how many times of) the height difference at which to cut the trees should be.
