## Supplemental Table S1 for "SHARP: Single-cell RNA-seq Hyper-fast and Accurate Processing via Ensemble Random Projection"

**Supplemental Table S1** scRNA-seq datasets used in this paper. The dataset of 1.3 million single cells does not provide the reference clusters so far. Please refer to the main manuscript for the references of all datasets.

| Dataset | Organism | No. of cells | No. of cluters |
|---|---|---|---|
| Goolam | Mouse | 124 | 5 |
| Darmanis | Human | 420 | 8 |
| Wang | Human | 479 | 8 |
| Kolod | Human | 704 | 3 |
| Baron_m1 | Mouse | 822 | 13 |
| Baron_m2 | Mouse | 1064 | 13 |
| Baron_h4 | Human | 1303 | 14 |
| Baron_h2 | Human | 1724 | 14 |
| Baron_h1 | Human | 1937 | 14 |
| Enge | Human | 2282 | 6 |
| Klein | Human | 2717 | 4 |
| Baron_h3 | Human | 3605 | 14 |
| Montoro_small | Mouse | 7193 | 7 |
| Park | Mouse | 43745 | 16 |
| Macosko | Mouse | 44808 | 39 |
| Montoro_large | Mouse | 66265 | 13 |
| 1.3 million | Mouse | 1306127 | NA* |
