## Supplemental Table S2 for "SHARP: Single-cell RNA-seq Hyper-fast and Accurate Processing via Ensemble Random Projection"

**Supplemental Table S2** Running time (in minutes) of SHARP against existing methods for the three simulated scRNA-seq datasets. Numbers in bracket are the number of cells in each dataset. $m \pm n$: mean ± standard deviation. SHARP, SC3, tSNE+kMeans and UMAP+kMeans use stochastic algorithms, thus mean ± standard deviation of the results are reported. Note that when the number of cells is larger than 5000, SIMLR uses smaller number of iterations to improved efficiency. The results are based on single-core measurement.

| Dataset / Method | **mdata3 (3898)** | **mdata6 (16717)** | **mdata8 (37538)** |
|---|---|---|---|
| SHARP | **3.22 ± 0.26** | **4.10 ± 0.15** | **9.25 ± 0.70** |
| SIMLR | 176.12 | 88.38 | 324.20 |
| SC3 | 13.33 ± 2.39 | 32.46 ± 2.99 | 45.64 ± 12.74 |
| hclust | 68.6 | 250.14 | 745.93 |
| tSNE-kMeans | 33.48 ± 18.47 | 403.87 ± 42.95 | 1186.91 ± 92.16 |
| UMAP-kMeans | 6.23 ± 0.11 | 131.28 ± 16.99 | 281.26 ± 37.72 |
| Seurat | 4.25 | 38.15 | 65.00 |
| CIDR | 5.85 | 218.08 | 2196.31 |
| Phenograph | 9.28 | 122.92 | 540.57 |
