## Supplemental Table S3 for "SHARP: Single-cell RNA-seq Hyper-fast and Accurate Processing via Ensemble Random Projection"

**Supplemental Table S3** Functional analysis for the 1.3 million single cells. Only those clusters with at least 1000 cells are listed here. See the paper for how to obtain the highly expressed genes. The pathway analysis is done by using Enrichr (Kuleshov et al. 2016). Typical functional terms are listed with p-value.

| Cluster No. | No. of cells | No. of highly expressed genes | Typical Functional Terms (p-value) |
|---|---|---|---|
| 1 | 390991 | 60 | positive regulation of fat cell apoptotic process (5e-4); neuron projection development (6e-4); neuron projection morphogenesis (7e-4). |
| 2 | 223822 | 692 | dendritic spine morphogenesis (2e-8); regulation of autophagosome assembly (3e-7); regulation of aggrephagy (7e-7). |
| 3 | 189734 | 436 | axon guidance (3e-9); synapse assembly (8e-8); chemoattraction of axon (1e-7). |
| 4 | 153088 | 83 | Rho protein signal transduction (7e-3); ARF protein signal transduction (2e-2); Ral protein signal transduction (2e-2) |
| 5 | 98089 | 5 | trigeminal nerve morphogenesis (1e-3); photoreceptor cell fate specification (1e-3); regulation of cell cycle phase transition (1e-3). |
| 6 | 75239 | 148 | calcium ion regulated exocytosis (2e-4); ephrin receptor signaling pathway (9e-4); dense core granule exocytosis (1e-3). |

**Supplemental Table S3** Functional analysis for the 1.3 million single cells. Only those clusters with at least 1000 cells are listed here. See the paper for how to obtain the highly expressed genes. The pathway analysis is done by using Enrichr (Kuleshov et al. 2016). Typical functional terms are listed with p-value.

| | | | |
|---|---|---|---|
| 7 | 70406 | 2267 | SRP-dependent cotranslational protein targeting to membrane (8e-13);<br>translational elongation (6e-11);<br>fatty acid beta-oxidation using acyl-CoA oxidase (1e-8). |
| 8 | 56749 | 1231 | DNA damage checkpoint (4e-8);<br>non-motile cilium assembly (2e-7);<br>regulation of lipid kinase activity (1e-6). |
| 9 | 23112 | 553 | negative regulation of antisense RNA transcription (5e-6);<br>nitrogen catabolite repression of transcription (5e-6);<br>carbon catabolite repression of transcription (5e-6). |
| 10 | 3610 | 1 | None |
| 11 | 3552 | 18 | establishment of melanosome localization (5e-3);<br>apical protein localization (5e-3);<br>ear development (5e-3). |
| 12 | 1608 | 2366 | Parkinson’s disease (1e-29);<br>mitochondrial translation (5e-26);<br>translational termination (4e-26). |
| 13 | 1526 | 0 | None |

**Supplemental Table S3** Functional analysis for the 1.3 million single cells. Only those clusters with at least 1000 cells are listed here. See the paper for how to obtain the highly expressed genes. The pathway analysis is done by using Enrichr (Kuleshov et al. 2016). Typical functional terms are listed with p-value.

| | | | |
|---|---|---|---|
| 14 | 1227 | 7 | positive regulation of interleukin-13 biosynthetic process (2e-3);<br>monocyte activation involved in immune response (2e-3);<br>regulation of isotype switching (3e-3). |
| 15 | 1213 | 2 | None |
| 16 | 1082 | 4 | None |
| 17 | 1051 | 5 | None |
